# Personalized phosphoproteomics establish mTORC1 as a regulator of exercise-induced insulin sensitization in human skeletal muscle

**DOI:** 10.64898/2026.08.27.747624

**Authors:** Magnus R. Leandersson, Hannah Huckstep, Johan D. Onslev, Nicolai S. Henriksen, Jonas M. Kristensen, Magnus N. Zankel, Rasmus Elmquist, Joachim T.E. Madsen, Jakob S. Jensen, Jesper B. Birk, Kim A. Sjøberg, TingTing Wang, Janni Petersen, Jerry R. Greenfield, David E. James, Esben Søndergaard, Rasmus Kjøbsted, Sean J. Humphrey, Jørgen F.P. Wojtaszewski

**Author notes:** Correspondence (S.J.H), (J.F.P.W).

## Abstract

Exercise enhances skeletal muscle insulin sensitivity, but the signaling mechanisms responsible are poorly understood. Understanding them may open new therapeutic avenues for individuals with limited exercise capacity. Here, we used rapamycin to inhibit mTORC1 in combination with exercise and insulin stimulation in healthy men. A single dose of rapamycin enhanced the insulin-sensitizing effect of exercise by 53% on average compared to placebo. Responses varied widely across individuals (-40% to 218%), and we leveraged this variance through personalized phosphoproteomics to map the mTORC1-dependent signaling network in skeletal muscle. This identified the protein kinase MKNK2 as a candidate downstream effector, which we then targeted for functional validation. Pharmacological inhibition of MKNK2 with eFT508 in insulin-clamped mice reduced both whole-body and skeletal muscle insulin sensitivity, confirming a functional role for MKNK2 activity in muscle glucose uptake. We then used eFT508 in *ex vivo* incubated human skeletal muscle to map the signaling network downstream of MKNK2, identifying the translational initiator eIF4G1 as a further regulatory node. Together, these findings indicate that exercise-induced insulin sensitization is actively constrained by a negative feedback pathway running from mTORC1 through the translational regulators MKNK2 and eIF4G1, raising the possibility that rapid translation of unidentified target proteins contributes to fine-tuning glucose uptake.

## Introduction

Insulin resistance is an underlying pathological condition driving the development of chronic diseases, including type 2 diabetes^1^, cardiovascular disease^2^, and several types of cancer^3–5^. Chronic hyperglycemia and the resulting compensatory hyperinsulinemia can both promote disease pathogenesis (reviewed in ^6^). While several classes of glucose-lowering pharmacotherapies exist, including GLP-1 receptor agonists, sulfonylureas, and metformin, these often prove insufficient to halt disease progression even as combination therapies^7,8^. Moreover, many of these agents primarily target glycemic control by elevating endogenous insulin secretion, which fails to resolve the underlying insulin resistance.

Conversely, both acute exercise^9^ and exercise training^10^ enhance whole-body insulin sensitivity, lowering the circulating insulin required to maintain glucose homeostasis^11,12^. Importantly, exercise-centered interventions outperform standard pharmacotherapy in preventing insulin resistance progression^13,14^. Because voluntary exercise is not always feasible or sustained, considerable interest has emerged in pharmacologically targeting this sensitizing effect, a concept referred to as ‘exercise mimetics’^15–18^.

The insulin-sensitizing effect of exercise is confined to previously contracting muscle^19^ and involves potentiation of insulin-induced microvascular perfusion^20^, translocation of the glucose transporter GLUT4 to the sarcolemma^21^, and glucose storage as glycogen^22^. Together these processes enhance glucose delivery, transport, and intracellular flux, but the intracellular signaling nodes that coordinate them remain incompletely defined.

Protein phosphorylation provides a rapid mechanism for cellular adaptation, and both insulin^23–27^ and exercise^24,28^ extensively remodel the skeletal muscle phosphoproteome. Phosphoproteomic analyses routinely identify thousands of regulatory sites, but connecting individual phosphorylation events to functional outcomes such as glucose uptake remains a challenge.

We recently developed an approach termed “personalized phosphoproteomics” that addresses this by exploiting phenotypic variance within a cohort to link individual phosphorylation events with physiological outcomes. Using this approach, we identified a strong association between mTORC1 signaling and exercise-induced insulin sensitization in young healthy men^29^, and recently replicated this finding in an independent cohort that included insulin-resistant individuals^23^. However, whether mTORC1 promotes or restrains exercise-induced insulin sensitization cannot be determined from associative data alone. Studies using the mTORC1 inhibitor rapamycin have implicated mTORC1 in reducing insulin sensitivity during hyperaminoacidemia in humans^30^, and following electrically stimulated contraction in rodents^31^, but under rested and unstimulated conditions rapamycin does not appear to alter insulin sensitivity^30–32^. Furthermore, if mTORC1 is a central orchestrator of exercise-induced insulin sensitization, the downstream effectors of mTORC1 in this context also remain largely undefined.

Here, we use rapamycin to test the causal role of mTORC1 in exercise-induced insulin sensitization in human skeletal muscle, and apply personalized phosphoproteomics to identify the downstream signaling network through which mTORC1 regulates this phenomenon.

## Results

### Rapamycin modulates whole-body metabolism and skeletal muscle insulin sensitivity

To explore the role of mTORC1 in muscle insulin sensitivity, we designed a crossover study where subjects received a single dose of rapamycin or placebo, followed by the assessment of leg glucose uptake prior to and during a hyperinsulinemic-euglycemic clamp. One hour of one-legged knee-extensor exercise was performed prior to the insulin clamp to induce an immediate enhancement of muscle insulin sensitivity in the previously exercised (PEX) leg compared to the non-exercised rested control (REST) leg. Hence, the experiment was internally controlled for both treatment (rapamycin versus placebo days) and activity (PEX versus REST legs) across exercise, exercise recovery, and insulin stimulation phases (**Figure 1A**). A total of 13 young and healthy males (**Table S1**) completed the study.

**Figure 1.**
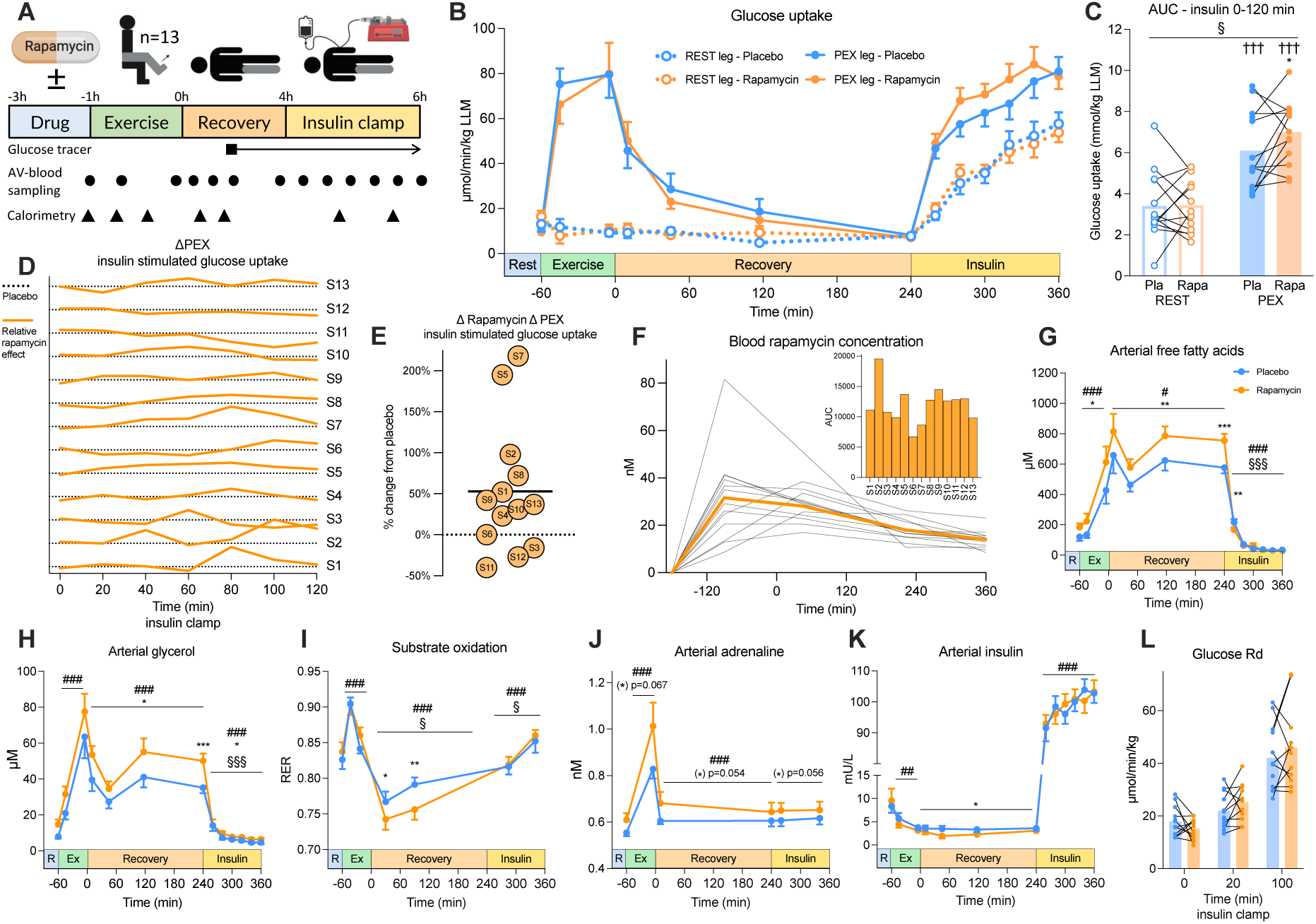
Rapamycin modulates whole-body metabolism and skeletal muscle insulin sensitivity. **A)** Experimental design and sampling timeline (n=13, unless otherwise stated). AV, arterio-venous. **B)** Leg glucose uptake throughout the study protocol (n=9 for the first 6 time points). LLM, lean leg mass; PEX, prior exercise. **C)** Area under the curve (AUC) for insulin-stimulated glucose uptake (0–120 min). **D)** Individual impact of rapamycin on exercise-induced insulin-stimulated glucose uptake. Each orange line represents a subject’s response on the rapamycin day, normalized to their own exercise effect on the placebo day (dotted horizontal baseline). **E)** Percentage change in exercise-induced insulin-stimulated glucose uptake following rapamycin treatment. **F)** Whole-blood rapamycin concentrations (grey: individual; orange: mean) and individual AUC. **G)** Arterial plasma free fatty acids and **H)** glycerol concentrations. **I)** Respiratory exchange ratio (RER), **J)** arterial plasma adrenaline, and **K)** arterial plasma insulin concentrations (n=9-13). **L)** Whole-body glucose turnover (Rd, rate of disappearance) during the insulin clamp (n=11-12). Data in **[B, G–K]** are means ± SEM; data in **[C, L]** are means with individual values paired (placebo vs. rapamycin). Two-way repeated measures ANOVA was used to test the following factors: *rapamycin* vs. *exercise* **[C]**; *time* vs. *rapamycin* within each condition (exercise, recovery, or insulin), using the final time point of the preceding condition as baseline **[G–K]**; and *rapamycin* vs. *time* **[L]**. Straight lines indicate a main effect. Significant interactions (p<0.05) were followed by Tukey’s post hoc tests. §§§ p<0.001, § p<0.05 *interaction* of factors; ††† p<0.001 effect of *exercise*; *** p<0.001, ** p<0.01, * p<0.05 effect of *rapamycin*; ### p<0.001, ## p<0.01, # p<0.05 effect of *time*.

As expected, on the placebo day, exercise acutely increased glucose uptake, which subsequently returned to baseline levels in exercise recovery. During subsequent insulin stimulation, the PEX leg demonstrated an enhanced sensitivity compared to the REST leg (**Figure 1B**). While rapamycin did not affect leg glucose uptake during exercise, during exercise recovery, or in the REST leg during insulin stimulation, the insulin-sensitizing effect in the PEX leg was significantly greater compared to the placebo control (**Figures 1B-C**). Notably, some subjects exhibited a greater effect of rapamycin on exercise-induced insulin sensitization during the early phase of insulin stimulation (S5, S10, S13), while others showed a more prominent effect during the late phase (S1, S2, S6, S7) (**Figure 1D**). We therefore quantified the overall effect of rapamycin by calculating the area under the curve (AUC) for the difference in glucose uptake between the PEX and REST leg over the 2h insulin stimulation period. This analysis demonstrated that, while prior exercise markedly enhanced muscle insulin sensitivity, administration of rapamycin further augmented this response, averaging a 53% improvement of the exercise effect (**Figure 1E**). These data extend our previous phosphoproteomic observations^23,29^ and are consistent with a model in which mTORC1 signaling acts through a negative feedback pathway to constrain muscle glucose uptake.

Rapamycin remained systemically available throughout the entire study day (**Figure 1F**) and broadly impacted whole-body metabolism. Plasma concentrations of free fatty acids and glycerol were elevated by rapamycin during most of the study protocol (**Figures 1G-H**), coinciding with a greater reliance on whole-body fat oxidation during early exercise recovery (**Figure 1I**). Circulating catecholamines tended to be higher following rapamycin administration (**Figures 1J** and **S1F**), which likely contributed to both the increased lipid availability (**Figures 1G-H**) and the slight reduction in plasma insulin concentrations prior to the clamp (**Figure 1K**). Importantly, steady-state insulin levels were unaffected by rapamycin during the hyperinsulinemic clamp (**Figure 1K**). Furthermore, the elevated lipid availability and catecholamine levels did not alter whole-body insulin sensitivity (**Figure 1L**). These data suggest that the enhanced insulin-sensitizing effect of exercise upon rapamycin administration is driven by skeletal muscle-intrinsic factors induced by the interaction of rapamycin, insulin, and exercise.

### Rapamycin reduces the rate of protein synthesis, but not protein breakdown, in skeletal muscle

Given the importance of mTORC1 in upholding muscle protein synthesis^33–38^, we hypothesized that the combination of prior exercise and insulin stimulation would significantly elevate the rate of protein synthesis in an mTORC1-dependent manner, as this combination has previously been shown to synergistically enhance mTORC1 signaling^23,29,39,40^. We thus implemented a continuous infusion of [^13^C_6_]-phenylalanine to assess the effect of rapamycin on protein synthesis and breakdown during exercise recovery, as well as during the subsequent insulin stimulation in a subset of the study group (n=7-9) (**Figure 2A**). In the exercise recovery period, rapamycin reduced the rate of muscle protein synthesis irrespective of PEX or REST status (**Figure 2B**). This aligns with previous observations in the recovery period from exercise^34–37^, though some report this effect to be specifically confined to PEX muscle^36,37,41^, which is likely dependent on the exercise modality and the circulating levels of amino acids.

**Figure 2.**
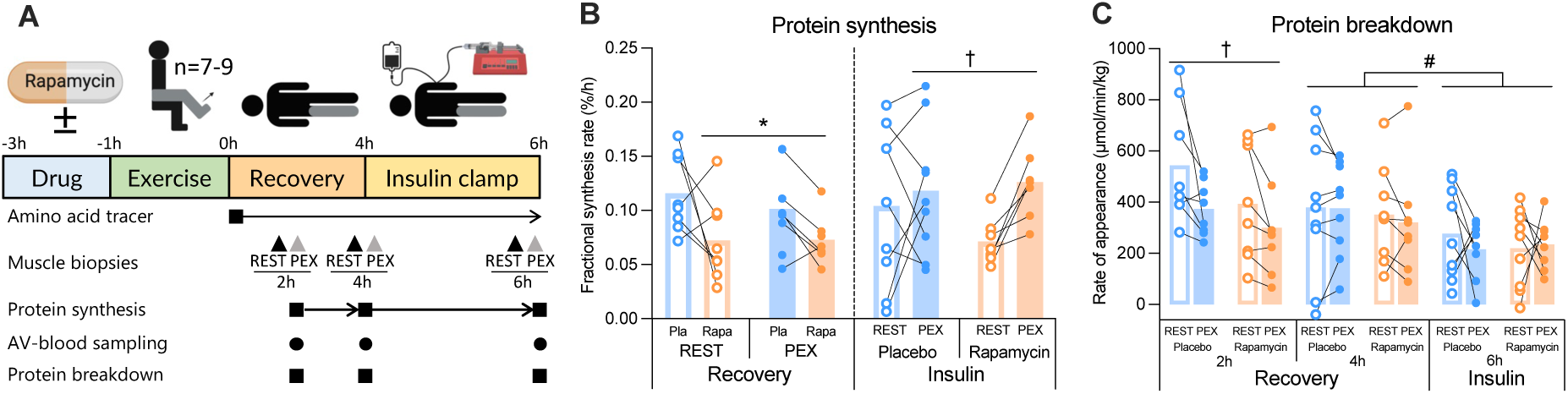
Rapamycin reduces skeletal muscle protein synthesis without affecting protein breakdown. **A)** Experimental timeline and sampling protocol for muscle protein kinetics (n=7-9). AV, arterio-venous; PEX, prior exercise. **B)** Skeletal muscle protein synthesis rates based on amino acid (AA) tracer incorporation during exercise recovery and insulin-stimulated periods following one-legged exercise. Note: For visual clarity, pairing of individual data points are flipped between exercise recovery and insulin stimulation to highlight the significant effect. **C)** Protein breakdown rates (rate of appearance, Ra) across the leg at specific time points following one-legged exercise, determined by AA tracer dilution. Data in **[B–C]** are presented as means with individual values paired across study days (placebo vs. rapamycin) or across legs (REST vs. PEX). Two-way repeated measures ANOVA was used to test the factors *rapamycin* vs. *exercise* at each time point independently **[B, C]**. Subsequently, all four groups were pooled at each time point to test the effect of *time* via paired t-test. Main effects are indicated by straight lines. † p<0.05 effect of *exercise*; * p<0.05 effect of *rapamycin*; # p<0.05 effect of *time*.

To our knowledge, no study has reported on the rate of muscle protein synthesis during insulin stimulation in combination with prior exercise (i.e., the condition where mTORC1 signaling is potentiated). Notably, during insulin stimulation, the rate of muscle protein synthesis was significantly greater in the PEX leg compared to the REST leg (**Figure 2B**). While a two-way ANOVA revealed a significant main effect of prior exercise, no significant interaction was observed (p = 0.21), precluding the statistical distinction of individual placebo and rapamycin effects. However, a closer inspection of the individual paired data (REST vs. PEX) indicates that this main effect was primarily driven by a consistent increase in the rapamycin condition, while no such trend was apparent with placebo (**Figure 2B**). This was contrary to our hypothesis. The blunted response with placebo may be related to the experimental conditions of the hyperinsulinemic-euglycemic clamp, which decreases plasma amino acid availability through the inhibition of whole-body protein breakdown, thereby potentially limiting anabolic signaling effects^42^. In contrast, the rapamycin condition exhibited a lower baseline rate of protein synthesis leading up to the clamp (i.e., during exercise recovery). Consequently, increasing the synthesis rate from this suppressed baseline may depend on factors other than amino acid availability. This unexpected but noteworthy observation suggests that the combination of prior exercise and insulin stimulation can rescue the rapamycin-induced suppression of protein synthesis.

Measurements of protein breakdown revealed a PEX-induced reduction at 2h into exercise recovery, as well as an insulin-induced reduction irrespective of prior exercise status (**Figure 2C**). No effects of rapamycin on protein breakdown were observed in any condition (**Figure 2C**).

These data suggest that the primary influence of mTORC1 on muscle protein kinetics is restricted to protein synthesis. Together with the systemic data in **Figure 1**, these results show that a single oral dose of rapamycin is sufficient to alter protein, lipid, and carbohydrate metabolism in humans, underscoring the broad metabolic role of mTORC1 signaling.

### Personalized phosphoproteomics identifies a subset of sites linked to rapamycin-induced improvements in muscle insulin sensitivity following exercise

Having established a causal link between mTORC1 signaling and muscle insulin sensitivity following exercise (**Figure 1C**), we next aimed to apply personalized phosphoproteomics to uncover the downstream targets of mTORC1 related to this rapamycin-induced phenotype (**Figure 3A**). To evaluate the pharmacological effect of rapamycin across different physiological phases, we defined four distinct analytical conditions from the muscle biopsies obtained pre- and post-insulin clamp (**Figure 3A**):

1. First, an exercise-primed state, representing the absolute phosphorylation status in the PEX leg prior to insulin stimulation (PEX basal).
2. Second, a baseline-adjusted exercise-primed state, calculated as the phosphorylation difference between the PEX and REST legs pre-insulin (ΔPEX basal).
3. Third, an insulin-responsive state, representing the absolute phosphorylation status in the PEX leg during the clamp (PEX insulin).
4. Finally, a baseline-adjusted insulin-responsive state, calculated as the phosphorylation difference between the PEX and REST legs during the clamp (ΔPEX insulin).

**Figure 3.**
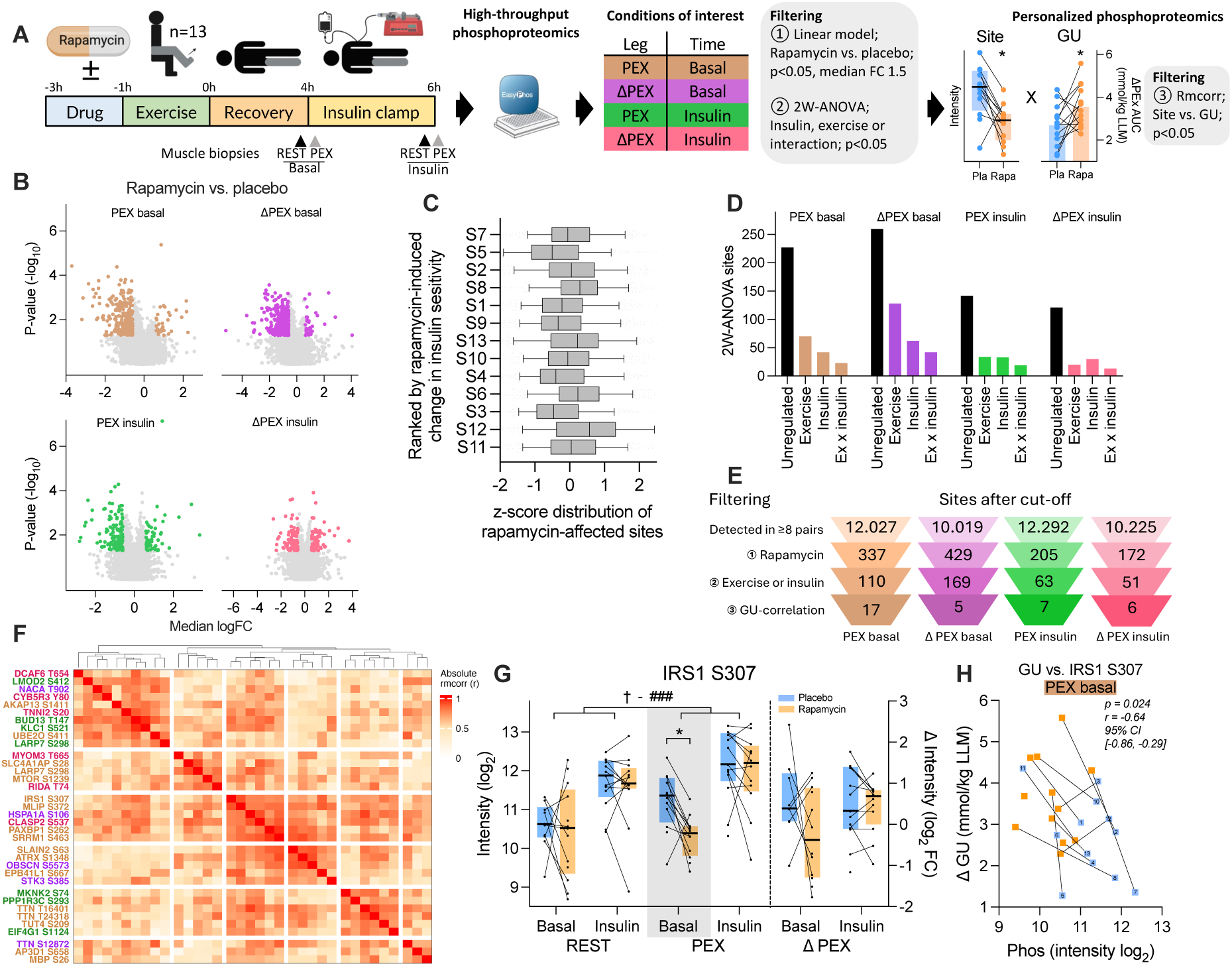
Personalized phosphoproteomics identify a subset of sites linked to rapamycin-induced improvements in muscle insulin sensitivity following exercise. **A)** Experimental timeline and 3-step statistical filtering workflow for personalized phosphoproteomics (n=13). AUC, area under the curve; GU, glucose uptake; PEX, prior exercise; Rmcorr, repeated measures correlation. **B)** Rapamycin-responsive phosphorylation sites (p < 0.05, median FC > 1.5) across defined skeletal muscle biopsy conditions. **C)** Distribution of z-scored rapamycin-responsive sites (from B). Sites with a median < 0 were inverted (multiplied by -1), and all values were z-score normalized across the cohort (i.e., the greatest rapamycin response corresponds to the highest z-score). Subjects are ranked by their rapamycin-induced change in insulin sensitivity (from Figure 1E), with the greatest responder at the top. **D)** Number of rapamycin-responsive sites regulated by insulin and/or exercise, determined by two-way repeated measures ANOVA (p < 0.05). **E)** Schematic of the 3-step statistical filtering process (as denoted in A). **F)** Rmcorr matrix of glucose uptake-correlated sites across all conditions. **G)** Phosphorylation levels and **H)** correlation with rapamycin-induced change in glucose uptake at PEX basal for IRS1 S307. Glucose uptake data is presented as the change in AUC between REST and PEX leg across the entire insulin clamp (0-120 min). Data in **[A (Site), G]** are presented as medians ± interquartile range; data in **[A (GU)]** are mean with individual values paired across study days (placebo vs. rapamycin). Data in **[H]** are individual paired values. Statistical analyses included: paired linear models to test the effect of *rapamycin* vs *placebo* [B, G]; two-way repeated measures ANOVA to test the factors *exercise* vs. *insulin*, independent of rapamycin treatment **[D, G]**; and Rmcorr to evaluate associations between glucose uptake and phosphorylation sites **[E, H]** or inter-site correlations **[F]**. Main effects are indicated by straight lines. * p<0.05 effect of *rapamycin*; † p<0.05 effect of *exercise*; ### p<0.001 effect of *insulin*.

Across the four defined conditions, rapamycin significantly affected the regulation (p<0.05, absolute median fold change > 1.5) of 172-429 phosphosites (**Figure 3B+E**), amounting to ∼2-4% of the quantified phosphoproteome. This relative modest scope of regulation contrasts with previous cell culture^43–45^ and rodent-based^46^ studies, which have reported that ∼8-20% of the phosphoproteome is regulated by rapamycin. However, these cell culture^43–45^ studies used *in vitro* incubation concentrations of 100-550 nM, while the rodent^46^ study used an intraperitoneal injection dose of rapamycin at 1.0 mg/kg bodyweight. In contrast, the maximum safe dose for a single oral ingestion of rapamycin in humans is 0.21 mg/kg bodyweight^47^, which was the dosage used in the present study. The circulating levels of rapamycin in whole blood ranged from ∼20 to 40 nM (**Figure 1F**). However, because rapamycin is highly sequestered in erythrocytes, exhibiting a 38:1 whole blood-to-plasma ratio^47^, the bioavailable plasma concentration of rapamycin was likely only in the range of ∼0.5-1.0 nM across the study day, with an estimated peak plasma exposure (*C_max_*) of ∼3 nM^47^. These pharmacokinetic data indicate relatively low tissue exposure of rapamycin in our human *in vivo* study in comparison to other models, which likely explains the milder effect on the global phosphoproteome. Notably, however, this dose was still sufficient to broadly impact metabolic regulation (**Figure 1** and **2**).

We observed substantial inter-individual variation in the rapamycin-induced improvement in muscle insulin sensitivity following exercise (ranging from -40% to 218%; **Figure 1E**), which could not be explained by differences in drug metabolism as the circulating rapamycin availability was generally similar across the cohort (**Figure 1F** and **S1G**). We therefore investigated whether this phenotypic variability could be explained by global differences in the rapamycin-affected phosphoproteome (i.e., testing whether high-responding subjects exhibited a more robust global intracellular response to rapamycin). Subjects 11 and 12 exhibited the weakest effects of rapamycin on muscle insulin sensitivity, whereas Subjects 5 and 7 exhibited the strongest effects (**Figure 1E**). However, upon inspecting the relative signaling responses to rapamycin (where a higher *z*-score represents a greater relative drug effect compared to the overall cohort), the *z*-score distribution of all rapamycin-affected sites was highest for Subject 12 and lowest for Subject 5 across the cohort (**Figure 3C**). There was no association between the rapamycin-induced changes in insulin sensitivity and global signalling (**Figure S2C+D**). These observations demonstrate that the insulin sensitivity phenotype was not driven by global differences in the intracellular pharmacological response, but rather indicate that a specific subset of phosphosites likely mediates the physiological effect.

The rapamycin effect on glucose uptake is superimposed on the combined effect of insulin and exercise (**Figure 1B**). Therefore, we next filtered for phosphosites regulated (p<0.05) by either exercise, insulin, or the interaction of these factors, ignoring the drug condition for this analytical step (**Figure 3D**). Subsequently, we correlated the phosphorylation of each site with the rapamycin-induced change in insulin sensitivity (*GU in* **Figure 3A** *derived from* **Figure 1C**), identifying a total of 36 sites across the four different conditions of interest (p<0.05 with a bootstrapped 95% CI excluding 0) (**Figure 3E+F**). Given the unadjusted thresholds applied at each filtering step, these sites should be considered hypothesis-generating candidates requiring independent validation.

### Re-evaluating the canonical mTORC1-S6K-IRS1 feedback pathway

Previous studies linking mTORC1 to the negative regulation of insulin sensitivity, primarily under hyperphysiological or *in vitro* conditions, have suggested a pathway involving an mTORC1-S6K-IRS1 axis leading to reduced proximal insulin signaling^30,31,48–52^. This pathway includes regulation of several inhibitory phosphorylation sites on IRS1, including S307, S312, and S636. Interestingly, in our analytical pipeline, IRS1 S307 met all three regulatory criteria: it was regulated by rapamycin in the PEX basal state, regulated by insulin and exercise, and correlated with the rapamycin-induced change in insulin sensitivity (**Figure 3G**+**H**). Supported by the previous reports on IRS1 regulation^30,31,48–52^, this initially points to IRS1 S307 as a prime candidate underlying the rapamycin-induced insulin sensitivity phenotype.

However, a closer inspection of the IRS1 S307 data in our setup revealed that the rapamycin effect at PEX basal was primarily driven by changes in the placebo group (**Figure 3G**). Specifically, we observed an increased phosphorylation in the PEX basal vs. REST basal state with placebo administration, while no such change was observed during rapamycin administration (**Figure 3G**). According to the proposed inhibitory role of IRS1 S307 phosphorylation, this would theoretically reduce insulin signaling in the PEX leg vs. the REST leg on the placebo day. However, regulation of the proximal insulin signaling node AKT was not different between PEX and REST, nor between rapamycin and placebo (**Figure S2E+F**). In fact, numerous studies^20,23,53–57^ support the consensus that proximal insulin signaling (from INSR to AKT) does not differ between PEX and REST under insulin-stimulated conditions.

Therefore, we interpret that the magnitude of increased IRS1 S307 phosphorylation in PEX basal vs. REST basal with placebo, and the reduced phosphorylation in rapamycin vs. placebo under PEX basal conditions, is not sufficiently strong to propagate downstream in our normo-physiological setup. At least not through the canonical insulin signaling axis. This does not exclude the possibility that the mTORC1-S6K-IRS1 pathway could impact canonical insulin signaling under different conditions or in different models. We thus deduce that the regulation of IRS1 S307 serves as a positive control from an mTORC1 signalling perspective, despite having negligible mechanistic relevance in this setup, and propose that other mTORC1-regulated phosphosites must drive the changes in exercise-induced muscle insulin sensitivity following rapamycin treatment.

### Disentangling the functional role of identified phosphosites in the rapamycin-induced enhancement of insulin-stimulated glucose uptake

The most robustly rapamycin-affected site across all conditions was mTOR S1239 (**Figure 4A**), an uncharacterized site that correlated significantly with the rapamycin-induced change in insulin sensitivity in the PEX basal state (**Figure 4B**). To investigate the functional relevance of this phosphorylation site, we substituted the orthologous residue in fission yeast (**Figure 4C**) with a non-phosphorylatable alanine (Tor1.S1097A). We compared survival of this mutant with that of wild-type (WT) and a previously established hyperactive mutant (Tor1.I1816T;^58^) in a glucose starvation assay. Notably, the Tor1.S1097A mutant exhibited significantly prolonged survival during glucose starvation, indicating reduced Tor1 activity when S1097 is mutated (**Figure 4C**). This is consistent with previous observations that rapamycin pretreatment, which reduces TORC1 activity in yeast, preserves viability during carbon starvation^59^. Notably, while the mTOR kinase can be incorporated into both mTOR complex 1 (mTORC1) and complex 2 (mTORC2), rapamycin specifically interacts with and inhibits mTORC1 in acute settings^60^. Similarly, during nutrient stress, fission yeast Tor1 can assemble into the orthologs of both human mTORC1 and mTORC2^61^. However, given the established rapamycin-specificity for mTORC1^60^, we interpret that the rapamycin-induced increase in mTOR S1239 phosphorylation observed in our human data likely occurs on mTOR exclusively within mTORC1. Extrapolating from the yeast data, where mutating Serine 1097 to alanine is consistent with reduced kinase activity, we infer that increasing mTOR S1239 phosphorylation in skeletal muscle could enhance mTORC1 activity, opposing the inhibitory effect of rapamycin. Thus, we speculate that the observed hyperphosphorylation of mTOR S1239 represents a compensatory feedback mechanism attempting to counter the rapamycin-induced inhibition. Whether phosphorylation at this specific site directly modulates the extent of rapamycin-induced mTORC1 inhibition in human skeletal muscle requires further investigation and should be the subject of future studies.

**Figure 4.**
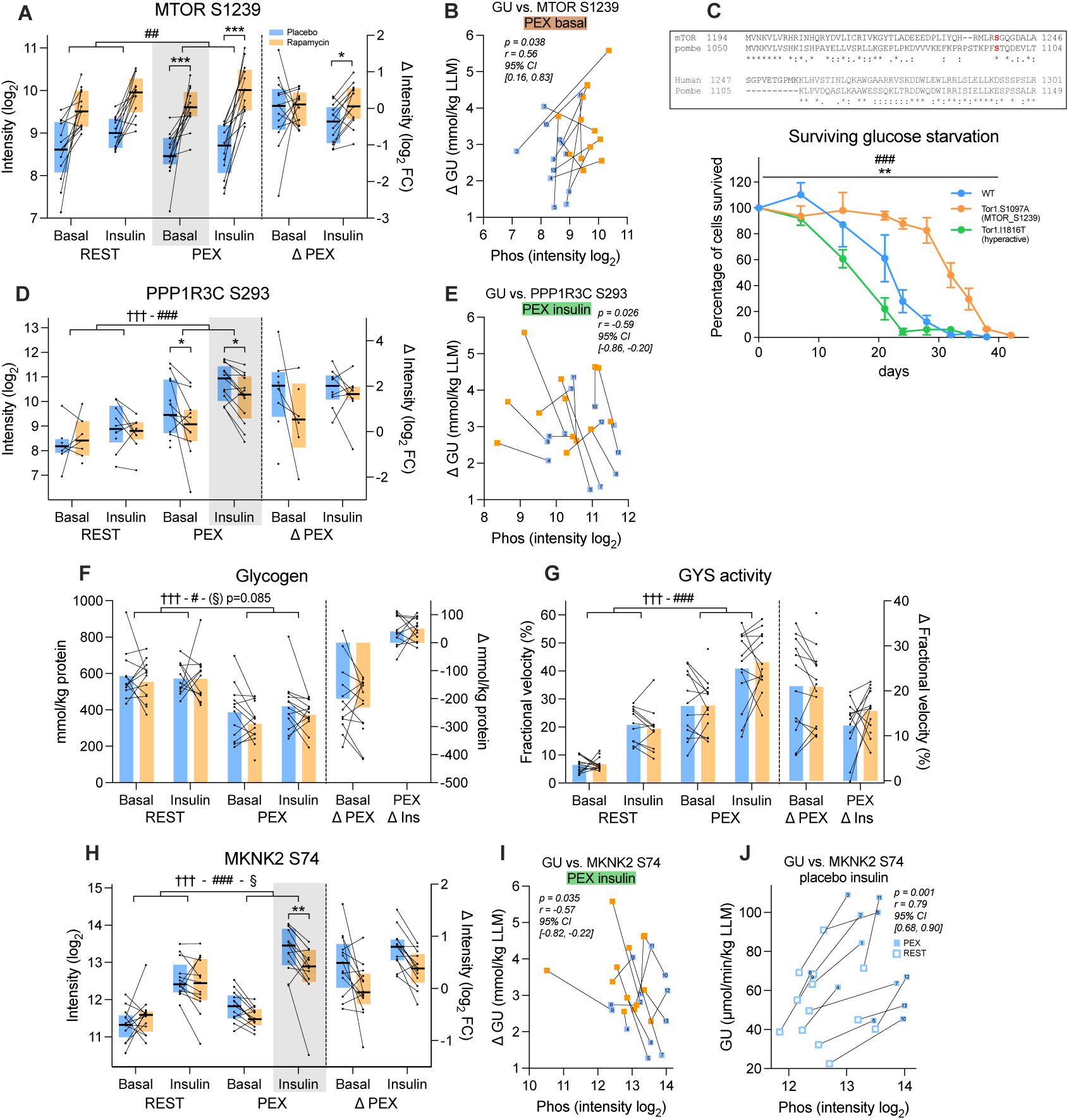
Disentangling the functional role of identified phosphorylation sites in rapamycin-induced change in insulin-stimulated glucose uptake. **A)** Phosphorylation levels and **B)** correlation with rapamycin-induced changes in glucose uptake for mTOR S1239 in human skeletal muscle. Glucose uptake data is presented as the change in AUC between REST and PEX leg across the entire insulin clamp (0-120 min). **C)** Cell survival assay during glucose starvation in fission yeast (*S. pombe*) expressing Tor1.S1097A (ortholog of human mTOR S1239), Tor1.I1816T (hyperactive control), or wild-type (WT). **D)** Phosphorylation levels and **E)** correlation with rapamycin-induced changes in glucose uptake for PPP1R3C S293. **F)** Muscle glycogen content and **G)** glycogen synthase activity. **H)** Phosphorylation levels of MKNK2 S74, with corresponding correlations between phosphorylation and **I)** rapamycin-induced or **J)** exercise-induced (placebo only) changes in glucose uptake. Glucose uptake (GU) data is presented as mean leg glucose uptake during the steady state of the insulin clamp (80-120 min). Data in **[A, D, H]** are medians ± interquartile range; data in **[B, E, I-J]** are individual paired values with subject IDs; data in [**C**] are means ± SEM; data in **[F-G]** are means with individual values. Lines connect samples from the same subject across study days (*placebo* vs. *rapamycin*) [**A-B, D-I**] or within study day (*REST* vs. *PEX*) [**J**]. Statistical analyses included: paired linear models to test the effect of *rapamycin* vs *placebo* [A, D, F-H]; two-way repeated measures ANOVA to test the factors *exercise* vs. *insulin*, independent of rapamycin treatment **[A, D, F-H]**; two-way mixed-design ANOVA (incorporating repeated measures for the *time* factor) was used to compare the factors *mutation (Tor1.S1097A* and *WT)* vs. *time* [C]; and Rmcorr to evaluate associations between glucose uptake and phosphorylation sites **[B, E, I-J]**. Main effects are indicated by straight lines. § p<0.05 *interaction* of factors; *** p<0.001, ** p<0.01, * p<0.05 effect of *rapamycin* or *Tor1.S1097A (vs. WT)*; ††† p<0.001 effect of *exercise*; ### p<0.001, ## p<0.01, # p<0.05 effect of *insulin* or time.

Another phospho-regulation of interest occurred on PPP1R3C S293. In the PEX insulin-stimulated state, phosphorylation of this site was significantly reduced by rapamycin. It was also upregulated by both exercise and insulin, and it correlated significantly with the rapamycin-induced change in insulin sensitivity (**Figure 4D**+**E**). PPP1R3C is one of seven known glycogen-targeting subunits that function to localize and actively regulate the catalytic activity of protein phosphatase 1 (PPP1CA)^62^. Notably, heterozygous PPP1R3C knockout (KO) mice exhibit reduced skeletal muscle glycogen levels, glycogen synthase (GYS) activity, and glucose uptake^63^. In our study, glycogen resynthesis in the PEX leg was modest during the relatively short 2-hour insulin clamp, with no observable differences between the rapamycin and placebo conditions (**Figure 4F**). However, this lack of divergence might be attributable to the low temporal resolution of this bulk tissue measurement. A potentially more sensitive molecular indicator of glucose flux into skeletal muscle is GYS activity. Yet, while GYS activity was dynamically regulated by both insulin and exercise, it was not impacted by rapamycin treatment (**Figure 4G**). PPP1R3C has previously been shown to bind several glycogen-related proteins in genetically modified cell models^62,64^. However, we were unable to sufficiently immunoprecipitate endogenous PPP1R3C from human muscle homogenates to determine whether rapamycin alters these interactions (data not shown). Future studies should explore the functional impact of PPP1R3C S293 phosphorylation, its direct relationship to glycogen metabolism, and whether this specific site contributes to the mTORC1-mediated negative feedback pathway on insulin signaling following exercise.

The phosphorylation of MKNK2 S74 was significantly regulated by the interaction between exercise and insulin, as well as by rapamycin in the PEX insulin-stimulated state (**Figure 4H**). Furthermore, the magnitude of this rapamycin-induced change in phosphorylation negatively correlated with the rapamycin-induced improvement in insulin sensitivity (**Figure 4I**). We have previously reported a strong positive correlation between MKNK2 S74 phosphorylation and the exercise-induced enhancement of insulin sensitivity in a cohort comprising both insulin-resistant and insulin-sensitive individuals^23^. We independently replicated this correlation in the current study using solely the data from the placebo day (**Figure 4J**). MKNK2 S74 is a known direct substrate of mTORC1, and phosphorylation at this specific site exerts an inhibitory effect on MKNK2 kinase activity^65^. Therefore, based on the synergistic increase in MKNK2 S74 phosphorylation driven by exercise and insulin (**Figure 4H**), its negative correlation with rapamycin-induced insulin sensitization (**Figure 4I**), and our replicated positive correlation with baseline exercise-induced sensitization (^23^ and **Figure 4J**), a potential mechanistic picture emerges: in the prior-exercised, insulin-stimulated state, MKNK2 activity is inhibited to restrain insulin sensitivity through an mTORC1-driven negative feedback pathway that is relieved by rapamycin.

### Pharmacological inhibition of MKNK2 reduces muscle insulin sensitivity

MKNK2 expresses a high baseline activity^66,67^ and is primarily regulated by inhibitory phosphorylation on S74^65^. Thus, the greatest inhibition of MKNK2 activity in the present study was driven by the combination of prior exercise and insulin stimulation on the placebo day (**Figure 4H**). To mimic this inhibitory effect and assess its impact on insulin sensitivity, we utilized the compound eFT508^68^ to pharmacologically inhibit MKNK2 *in vivo* in mice. Following an intravenous bolus infusion of eFT508 or a vehicle control, the mice underwent a 90-minute hyperinsulinemic-euglycemic clamp (**Figure 5A**). The injection of eFT508 significantly elevated blood glucose levels prior to the initiation of the clamp (**Figure 5B**-**C**), indicating an acute effect of MKNK2 inhibition on either endogenous glucose production or clearance. Importantly, however, the mice were subsequently clamped at identical blood glucose levels under hyperinsulinemic conditions, allowing us to exclusively evaluate peripheral insulin sensitivity (**Figure 5B**). The steady-state glucose infusion rate required to maintain euglycemia was significantly reduced following eFT508 administration (**Figure 5D**-**E**), demonstrating that the inhibition of MKNK2 impairs whole-body insulin sensitivity.

**Figure 5.**
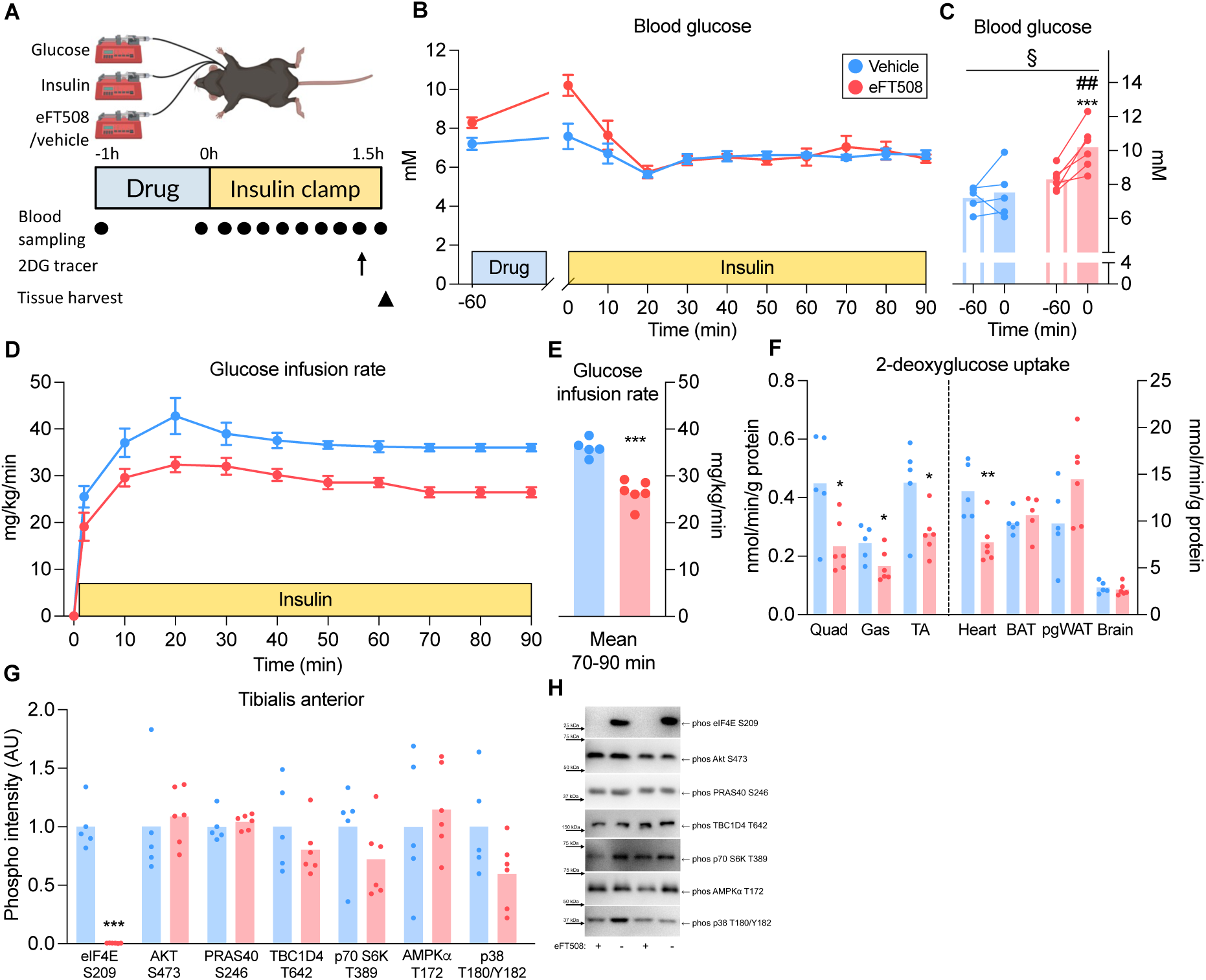
Pharmacological inhibition of MKNK2 reduces muscle insulin sensitivity. **A)** Experimental design and sampling timeline for acute eFT508 treatment in mice (n=5-6). 2DG, 2-deoxyglucose. **B)** Tail-vein blood glucose concentrations throughout the study protocol. **C)** Blood glucose levels before (–60 min) and after (0 min) intravenous bolus infusion of eFT508 or vehicle. **D)** Glucose infusion rate (GIR) during the hyperinsulinemic-euglycemic clamp. **E)** Mean GIR during the clamp steady state (70–90 min). **F)** Insulin-stimulated 2-deoxyglucose uptake in specific tissues during the steady state (80–90 min). Quad, Quadriceps; Gas, Gastrocnemius; TA, Tibialis Anterior; BAT, brown adipose tissue; pgWAT, perigonadal white adipose tissue. **G)** Immunoblotting of selective phosphorylation sites in the *tibialis anterior* muscle, and **H)** corresponding representative immunoblots. Data in **[B, D]** are means ± SEM; data in **[C, E–F]** are means with individual values. Lines connect samples from the same animal **[C]**. Statistical analyses included: Two-way ANOVA paired within drug group to test the factors *eFT508* vs. *time* **[C]**; and unpaired t-tests to compare *eFT508* vs. *vehicle* **[E–G]**. Significant interactions (p<0.05) were followed by Tukey’s post hoc tests. § p<0.05 *interaction* of factors; *** p<0.001, ** p<0.01, * p<0.05 effect of *eFT508*; ## p<0.01 effect of *time*.

During the final stage of the clamp, we infused a 2-deoxyglucose tracer to assess tissue-specific glucose uptake under steady-state conditions. Notably, this revealed that the eFT508-induced reduction in insulin sensitivity was confined to muscle tissue, including three distinct types of skeletal muscle and in cardiac muscle, whereas glucose uptake in the brain, as well as in brown and white adipose tissues, remained unaffected (**Figure 5F**). Immunoblotting confirmed the full inhibition of MKNK2 signaling, evidenced by the ablation of phosphorylation on the canonical downstream target, eIF4E S209. Conversely, canonical insulin, mTORC1, and stress-related signaling pathways remained generally unaltered (**Figure 5G-H+S3D-G**). These *in vivo* data support a role for MKNK2 in the regulation of muscle insulin sensitivity.

### Delineating the MKNK2 signaling axis in human skeletal muscle

While the upstream regulation of MKNK2 is well described^65,66,69,70^, its downstream signaling network remains largely unexplored, with only a single target site currently well-established in the literature (eIF4E S209)^67^. This specific site was not detected in our phosphoproteomic dataset, however targeted immunoblotting demonstrated increased phosphorylation following rapamycin treatment in the PEX leg under both basal and insulin-stimulated conditions (**Figure 6A-B**). This confirms the expected higher activity of MKNK2 following rapamycin treatment (**Figure 4H**). We next sought to identify the MKNK2 signalling network in human muscle. We thus combined a human *ex vivo* (HEV) muscle incubation model^71^ with pharmacological inhibition of MKNK2 signaling using eFT508 (**Figure 6C**). Because rapamycin prevents the inhibitory phosphorylation of MKNK2 S74 (**Figure 4H**), theoretically maintaining the kinase in a more active state, we hypothesized that sites sensitive to eFT508 inhibition should display regulatory patterns directly opposite to those induced by rapamycin. This pattern was demonstrated through targeted immunoblotting of the canonical eIF4E S209 (**Figure 6A** and **S4C**).

**Figure 6.**
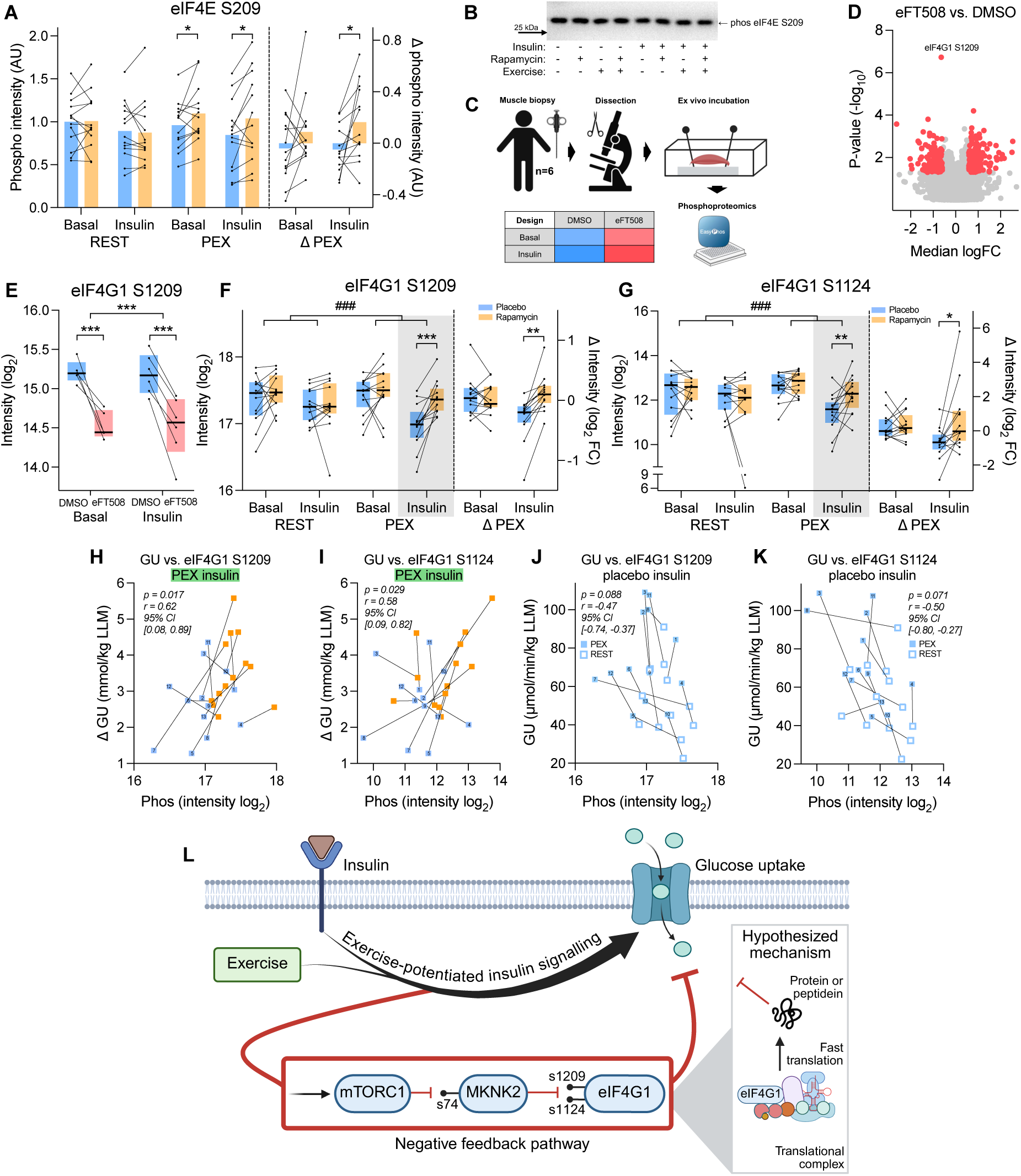
Delineating the MKNK2 signaling axis in human skeletal muscle. **A)** Immunoblotting of eIF4E S209 in skeletal muscle from human *in vivo* experiment, and **B)** corresponding representative immunoblots. **C)** Experimental design for *ex vivo* incubation of human skeletal muscle. **D)** Differentially phosphorylated sites responsive to eFT508 treatment (p<0.05, median FC > 1.5) in *ex vivo* muscle. **E)** Phosphorylation levels of eIF4G1 S1209 in human muscle incubated *ex vivo* with the MKNK2 inhibitor eFT508. Phosphorylation levels of **F)** eIF4G1 S1209 and **G)** eIF4G1 S1124 from human *in vivo* experiment, with corresponding correlations between phosphorylation and **H–I)** rapamycin-induced or **J–K)** exercise-induced (placebo only) changes in glucose uptake. Delta glucose uptake (ΔGU) data is presented as the change in AUC between REST and PEX leg across the entire insulin clamp (0-120 min). Glucose uptake (GU) data is presented as mean leg glucose uptake during the steady state of the insulin clamp (80-120 min). **L)** Schematic model illustrating the signaling pathways identified in the current study and the proposed downstream regulatory mechanisms. Data in **[A, E-G]** are presented as medians ± interquartile range; data in **[H-K]** are individual paired values with subject IDs; Lines connect samples from the same subject across drug condition (*placebo/DMSO* vs. *rapamycin/eFT508*) [**A, E-I**] or across legs (*REST* vs. *PEX*) [**J-K**]. Statistical analyses included: paired t-test [**A**] or linear models [**D-G**] to test the effect of drug treatment (*rapamycin/eFT508* vs *placebo/DMSO*); two-way repeated measures ANOVA to test the factors *exercise* vs. *insulin*, independent of rapamycin treatment **[A, F-G]**; and Rmcorr to evaluate associations between glucose uptake and phosphorylation sites **[H-K]**. Main effects are indicated by straight lines. *** p<0.001, ** p<0.01, * p<0.05 effect of *rapamycin* or *eFT508*; ### p<0.001 effect of *insulin*.

Surprisingly, phosphoproteomics analysis revealed that only a single phosphosite was robustly altered by eFT508 treatment (notably, eIF4E S209 was not detected) (**Figure 6D**). This site, eIF4G1 S1209 (**Figure 6E**), was not identified by the personalized phosphoproteomics pipeline, as it fell below the median fold-change threshold applied in the initial rapamycin analysis (FC = 1.33 versus a 1.5 cutoff) (**Figure 3B**). Guided by its emergence as the top eFT508-responsive site in the independent *ex vivo* experiment, we performed a hypothesis-driven re-analysis of eIF4G1 S1209 in the *in vivo* human phosphoproteomics dataset. This revealed a highly significant rapamycin effect, acting in the exact opposite direction of eFT508, specifically in the PEX insulin-stimulated state (p < 0.001) (**Figure 6F**). A related site on the same protein, eIF4G1 S1124, was successfully identified among the global rapamycin-affected sites (**Figure 3B**+**6G**) but, conversely, was not detected in the *ex vivo* human muscle setup. The regulation of eIF4G1 S1124 downstream of MKNK2 could therefore not be confirmed. Notably, however, the magnitude of the rapamycin effect on eIF4G1 S1124 strongly correlated with its effect on MKNK2 S74 (**Figure 3F**), indicating probable co-regulation.

Both sites (**Figure 6F-G**) display a regulatory pattern perfectly mirroring the expected downstream consequence of MKNK2 S74 inhibitory phosphorylation in response to both insulin and rapamycin (**Figure 4H**). More importantly, the phosphorylation of both eIF4G1 S1209 and eIF4G1 S1124 associated significantly with the rapamycin-induced change in insulin sensitivity (**Figure 6H-I**). We have previously reported that S1209 also correlates with the exercise-induced change in insulin sensitivity^23^. Consistent with this, in the present study, both sites exhibited a borderline association with the exercise-induced change in insulin sensitivity on the placebo day (**Figure 6J-K**).

While eIF4G1 S1209 was the only site robustly affected by eFT508 incubation (**Figure 6D**), we also assessed moderately but significantly affected sites using a cutoff similar to the one applied for rapamycin (p < 0.05, median FC > 1.5). We filtered for phosphosites regulated in the opposite direction by rapamycin (p < 0.05, median FC > 1.5) (**Figure 3B**) and manually evaluated the resulting 17 sites, identifying 10 with convincing regulatory patterns (**Figure S5A-T**). These 10 sites were located on proteins associated with cytoskeletal organization (CAVIN4, SYNE2), vesicle trafficking (RIN1, RAB40C), and gene expression and RNA processing (FAM120A, eIF3K, HNRNPD, RBMX, ZNF512B). While noteworthy, these 10 sites require additional experimental evidence to firmly establish them as part of the MKNK2 signaling network.

Collectively, these findings suggest that the mechanistic link between mTORC1 signaling and post-exercise insulin sensitivity^23,29^ operates through MKNK2 and extends downstream to eIF4G1. This signaling axis is now supported by the *in vivo* and *ex vivo* pharmacological manipulation of both mTORC1 and MKNK2. As previously established, the scaffolding protein eIF4G1 is required for canonical protein translation through its role in assembling the eIF4F initiation complex^72–76^. This suggests that the mTORC1-driven negative feedback pathway restraining muscle insulin sensitivity following exercise may operate through a rapid, translation-dependent response (**Figure 6L**).

## Discussion

In the present study, we utilized the kinase inhibitor rapamycin to demonstrate a central role for mTORC1 signaling in the regulation of human skeletal muscle insulin sensitivity during the post-exercise recovery period, acting as a negative feedback pathway to limit the rising influx of glucose.

The study builds upon our previous observations linking mTORC1 signaling to exercise-induced insulin sensitization in human skeletal muscle^23,29^, identified through a computational framework we term "personalized phosphoproteomics". Within the field of skeletal muscle physiology, associating omics-based molecular profiling with physiological phenotypes is an emerging experimental strategy^23,25,29,77–83^. However, a frequently overlooked aspect of these study designs is the tissue-specific resolution of the measured physiological phenotype. We view this as a continuum with inherent trade-offs: deeper tissue-specific physiological resolution typically necessitates a reduction in cohort size. Notably, our assessment of muscle insulin sensitivity, quantified via Fick’s principle-based leg glucose uptake (see *Methods*; REFs^23,29^), provides a much higher tissue-specific resolution than omics studies relying on systemic proxies for peripheral insulin sensitivity^21,74,81^. Conversely, those studies have many-fold larger cohorts, affording them greater statistical power. Which of these strategies proves more effective at identifying replicable molecular regulators of muscle insulin sensitivity remains to be seen. By demonstrating a causal link from mTORC1 signaling to muscle insulin sensitivity, the present study provides robust empirical support for the deep-phenotyping approach^23,29^. This mechanistic insight was generated despite two major pharmacological limitations associated with the *in vivo* use of rapamycin. First, the maximum safe oral dose in humans is restricted to 0.21 mg/kg^47^, which exerted only a submaximal inhibitory effect (see *Results*). Second, the allosteric nature of rapamycin only inhibits approximately 50% of all mTORC1 substrates, classically dividing the downstream signaling network into rapamycin-sensitive and rapamycin-resistant branches^84^. We therefore speculate that a more potent, global inhibition of mTORC1 signaling could yield an even greater impact on muscle insulin sensitivity.

Applying a three-step personalized phosphoproteomic pipeline (**Figure 3A**), we highlighted 36 candidate phosphosites associated with muscle insulin sensitivity. Several of these phosphosites may be causally linked to the regulation of glucose uptake through distinctive branching pathways. We explored the inhibitory phosphosite MKNK2 S74 in targeted follow-up studies, revealing a causal link between MKNK2 and muscle insulin sensitivity following the acute administration of the MKNK2 inhibitor eFT508 (**Figure 5F**). Interestingly, while our acute pharmacological inhibition of MKNK2 reduced both whole-body and muscle-specific insulin sensitivity, the chronic loss of MKNK2 via transgenic whole-body KO has been reported to protect against high-fat diet-induced insulin resistance^85^. Given that MKNK2 is a central regulator of translation initiation^66,67^, such divergent phenotypic outcomes between acute kinase inhibition and chronic genetic ablation are plausible due to long-term translational reprogramming or developmental compensation. Therefore, the careful selection of experimental models is important for disentangling the temporal role of MKNK2 in regulating muscle insulin sensitivity. Future investigations should employ structurally distinct pharmacological inhibitors alongside adult-onset inducible MKNK2 KO models to parse out primary signaling effects from secondary compensatory adaptations.

To explore the downstream signaling targets of MKNK2, we modified a classical HEV muscle incubation model to directly assess the effects of eFT508 treatment on human skeletal muscle. This modified HEV model was selected to maximize sequence coverage of the human skeletal muscle phosphoproteome, enabling a direct, tissue-matched comparison with our primary *in vivo* clinical data. The classical HEV model relies on surgical clamp-based open muscle biopsies^71^, a highly invasive and impractical procedure. In contrast, our modified HEV model utilizes muscle fiber bundles dissected from standard Bergström needle biopsies, a much less invasive approach that permits multiple consecutive samplings from the same subject. While the classical HEV model has been extensively validated^71,86–89^, the physiological fidelity of our modified version warrants further systematic characterization. However, for the present application, the physiological relevance of our model was supported by robust, dynamic phospho responses to titrations of both eFT508 (**Figure S4A**) and insulin (**Figure S4B**), as well as insulin-responsive GLUT4 translocation^90^. The present HEV experiment revealed that eIF4G1 S1209 is highly responsive to eFT508 (**Figure 6E**) and is regulated in the exact opposite direction by rapamycin (**Figure 6F**), strongly indicating that it is a downstream target of MKNK2. Surprisingly, eIF4G1 S1209 was the only phosphosite robustly displaying this divergent regulatory signature in the *ex vivo* phosphoproteomic dataset (notably, eIF4G1 S1124 and the canonical eIF4E S209 were not detected). We cannot exclude the possibility that other quantified phosphosites could be highly responsive to MKNK2 inhibition under different study designs or in different experimental models.

Ten additional phosphosites were considered probable MKNK2 downstream targets, based on their opposing responses to rapamycin, but these warrant further investigation to be firmly established (**Figure S5A-T**). Notably, 9 of these 10 sites presented with increased phosphorylation following eFT508 treatment, indicating the involvement of at least one additional enzyme between MKNK2 and these sites. The relatively low number of phosphosites downstream of MKNK2 is notable in comparison to other protein kinases (reviewed in ^91^), but may reflect a highly specialized substrate profile for MKNK2 or incomplete phosphopeptide coverage of the relevant target sites, as exemplified by the absence of the canonical substrate eIF4E S209 from the mass spectrometry data, despite confirmed regulation by immunoblotting (**Figure 6A**+**S4A**). A narrow target profile may also explain why, despite nearly 30 years since its discovery^92,93^, only one site has been firmly established as a direct downstream target of MKNK1 and MKNK2 (i.e., eIF4E S209).

While multiple distinct pathways concurrently regulate muscle insulin sensitivity following exercise, our analyses highlight the mTORC1-MKNK2-eIF4G1 signaling axis as a candidate mediator of the negative feedback pathway acting to restrain glucose uptake. This proposed axis is supported by targeted pharmacological inhibition and by the robust statistical associations between the phosphorylation status of MKNK2 S74, eIF4G1 S1209, and eIF4G1 S1124 and both the rapamycin-induced and exercise-induced changes in insulin sensitivity. Based on the established biological roles of MKNK2^66,67^ and eIF4G1^72–76^, our signaling data implicate acute translational regulation as part of this negative feedback pathway. We acknowledge that this interpretation is based on the established roles of these signaling components rather than direct measurements of translational output, which remain an important objective for future studies. This output may be in the form of known proteins or the newly described peptideins^94^, acutely functioning to modulate glucose uptake.

We note that a related hypothesis was tested previously, yielding evidence against the necessity of de novo protein synthesis for the acute regulation of muscle insulin sensitivity following a single bout of exercise^95^. That study, performed in rats, involved the surgical excision of the epitrochlearis muscle immediately post-exercise, followed by a 3.5-hour *ex vivo* incubation with the translation inhibitor cycloheximide prior to insulin stimulation, which reportedly did not impair the exercise-induced enhancement of insulin sensitivity. However, crucially, amino acids were absent from the incubation media throughout this prolonged *ex vivo* period. Because extracellular amino acid availability is required for the mTORC1-dependent protein synthesis^33,38,96^, the baseline translational capacity of the control group in that experimental setup was likely already suppressed. The translatability of those *ex vivo* rat findings to *in vivo* human physiology should therefore be reconsidered and a physiologically intact assessment of the role of mTORC1-mediated protein synthesis in the acute regulation of human muscle insulin sensitivity is warranted. While our phosphoproteomic data could theoretically accommodate a secondary, translation-independent role for eIF4G1, we propose that exercise-induced insulin sensitization in human skeletal muscle is restrained by an mTORC1-MKNK2-eIF4G1 negative feedback pathway, which may rely on the rapid translation of yet-unidentified target proteins or peptideins to fine-tune glucose uptake.

## Limitations

Our human studies only included healthy males of European ancestry within a narrow age range. In contrast, the mouse experiment only included female mice of the inbred C57BL/6J strain. While this demonstrates cross-sex conservation of the pathway across species, future studies should systematically include both sexes across all experiments and broaden human ancestry and mouse strains to improve the generalizability of our findings.

The administered dose of rapamycin was only sufficient to partially inhibit mTORC1 signaling (as discussed in *Results*), and we therefore used non-adjusted p-values in the bioinformatic pipeline. Our results may therefore include false positives, despite a three-step statistical filtering pipeline. The highlighted sites should thus be considered hypothesis-generating candidates requiring independent validation.

Our study extensively uses eFT508 to inhibit MKNK2 signaling. It is worth noting that eFT508 inhibits both MKNK1 and MKNK2, and the relative contribution of each isoform to the observed phenotype therefore cannot be distinguished from the present data. However, several lines of evidence suggest MKNK2 is the more relevant isoform in this context. MKNK2 is the predominantly expressed isoform in skeletal muscle, exhibits high constitutive activity^66^, and uniquely inhibits eIF4G1 S1147 phosphorylation in muscle through a mechanism not shared by MKNK1^97^. Moreover, basal eIF4E phosphorylation is reduced in Mknk2 but not Mknk1 KO mice^67^, consistent with a dominant role for MKNK2 in maintaining tonic signaling through this pathway in muscle.

It is also important to note that mice were studied at rest rather than following exercise. This study design was chosen to deconstruct the mTORC1-driven negative feedback pathway. Notably, while the activity of mTORC1 is potentiated by insulin stimulation following exercise^29,40^, MKNK2 activity, independent of mTORC1 regulation at S74, may not be dependent on exercise. We thus mimicked the exercise-induced mTORC1-driven inhibition of MKNK2 at S74 by simply studying pharmacological inhibition of MKNK2 at rest. Although we observed a ∼40% reduction in insulin stimulated glucose uptake into skeletal muscle (**Figure 5F**), this relative reduction may be more modest if studied in the recovery phase from exercise, where the negative feedback pathway is expected to signal in the placebo control group.

Finally, while we aimed to use the lowest dose of eFT508 that yielded a maximal response (**Figure S3B**+**S4B**), we cannot exclude the possibility that off-target effects could have an effect on either protein signaling in the HEV experiment and/or glucose clearance during the mouse clamp.

## Method

### Human in vivo experimental study

This study was approved by the Committee on Health Research Ethics of the Capital Region of Denmark (Reference number: H-20071202) and performed in accordance with the ethical standards laid down in the Declaration of Helsinki. All subjects provided written informed consent after receiving comprehensive oral and written information regarding the study procedures. The study was registered at ClinicalTrials.gov (identifier: NCT05233722).

#### Screening

Healthy male subjects were screened for study eligibility based on the following inclusion criteria: normal hematological parameters, no family history of diabetes or other known metabolic diseases, a normal body mass index (BMI; 18–25 kg/m^2^), age between 22 and 35 years, moderate physical activity levels (VO_2_ max: 40–60 mL/kg/min), and no current use of nicotine products. Initially, 40 potential participants were screened via telephone interviews, from which 25 proceeded to in-person clinical screening. Ultimately, 16 subjects met all criteria and were enrolled in the study. Three subjects withdrew prior to completing the experimental protocol; thus, a final cohort of 13 subjects (IDs: S1–S13) completed the entire study.

On two separate screening days, participants underwent a series of baseline assessments. Body composition was determined using dual-energy X-ray absorptiometry (DXA; DPX-IQ Lunar, Lunar Corporation), and maximal oxygen uptake (VO_2_ max) was assessed via indirect calorimetry (Vyntus CPX, Vyaire Medical) during an incremental cycling test. Subjects were then familiarized with the one-legged knee extensor ergometer^98^, with the designated exercising leg randomized (dominat vs. non-dominant) across participants. Finally, subjects performed an incremental one-legged knee extensor test to establish the peak workload (PWL) of the exercising leg. The PWL was defined as the final incremental step before oxygen uptake, carbon dioxide production, and heart rate deviated from a linear progression; a deviation from this linearity indicated the unintended recruitment of accessory muscle groups, thereby marking the physiological limit of the target muscle.

#### Experimental design

Three days prior to the first experimental study day, subjects were instructed to record their dietary intake, and they were asked to refrain from physical activity and alcohol consumption for two days prior to the experiment. On the morning of the experimental day, subjects arrived at the laboratory at 06:30 following an overnight fast. Subjects ingested a light, standardized breakfast consisting of oatmeal, skimmed or oat milk, and a small spoonful of sugar, which was estimated to constitute 5% of their daily energy intake. Next, subjects were administered either 0.21 mg/kg rapamycin (2 mg Sirolimus/Rapamune tablets; Pfizer) or 800 mg of calcium tablets as a placebo. The administration of either rapamycin or placebo on the first study day was randomized and executed by an unblinded clinical investigator; the subjects and all remaining experimental personnel were strictly blinded to the treatment allocation. Following drug ingestion, catheters (Pediatric Jugular Catheterization set; Arrow International) were inserted into both femoral veins and the femoral artery of the non-exercising leg under local anesthesia (∼3-5 mL of 10 mg/mL lidocaine without adrenaline [Xylocaine]; AstraZeneca).

At 08:30, subjects performed a one-hour bout of one-legged knee extensor exercise at 80% of their predetermined PWL, interspersed with two 5-minute intervals at 90% PWL. Following the exercise bout, subjects were returned to a hospital bed and rested for the remainder of the study day. To assess glucose kinetics, a stable glucose tracer ([6,6-²H₂]-glucose; Cambridge Isotope Laboratories) was infused through an antecubital venous catheter. The infusion was initiated two hours into the resting period with a bolus (14.3 µmol/kg) followed by a constant infusion (0.24 µmol/kg/min) to achieve steady-state conditions prior to the insulin clamp. For a subset of the subjects (S1–S9), a phenylalanine tracer (L-[ring-^13^C_6_]-phenylalanine) was initiated immediately following the exercise bout with a bolus (3 µmol/kg) followed by a constant infusion (0.05 µmol/kg/min).

Four hours post-exercise, a two-hour hyperinsulinemic-euglycemic clamp was initiated. Insulin (Actrapid; Novo Nordisk) was infused with an initial bolus of 9 mU/kg followed by a constant rate of 1.42 mU/kg/min. This infusion rate targets a steady-state plasma insulin concentration of ∼100 mU/L, mimicking high-physiological postprandial levels observed after a mixed meal^39^. Concurrently, exogenous glucose infusion was adjusted every ∼5 minutes to maintain euglycemia. The exogenous glucose was supplied from a 10% glucose infusion bag (Fresenius Kabi) artificially enriched with 1.9% [6,6-^2^H_2_]-glucose to maintain steady-state tracer enrichment levels throughout the clamp.

Four weeks after the first experimental day, subjects returned for their second experimental visit. Subjects were instructed to strictly replicate their recorded food intake from the three days leading up to the first visit. An identical experimental protocol was conducted, with the sole exception that subjects received the opposite treatment (rapamycin or placebo) in accordance with the randomized crossover design.

#### Experimental measurements

During the experimental study days, a rigorous sampling protocol was executed to obtain blood, blood flow, muscle tissue, and respiratory data.

*Blood samples* were drawn from the three femoral catheters (one arterial, two venous) at 13 distinct time points throughout the study day (**Figure 1A**). Samples were collected using syringes with or without heparin coating. Aliquots of heparinized whole blood were immediately analyzed using a blood gas analyzer (ABL800 FLEX; Radiometer) to determine glucose concentrations. Other blood parameters measured simultaneously (including lactate, electrolytes, and gases) were used solely for clinical monitoring and are not reported. The majority of the remaining blood aliquots were immediately centrifuged to isolate plasma, which was subsequently frozen on dry ice. Selected non-heparinized arterial samples were collected in tubes containing trace amounts of EDTA and stored as whole blood. Additionally, arterial blood was drawn into heparinized syringes every ∼5 minutes during the hyperinsulinemic-euglycemic clamp for immediate blood glucose analysis (ABL800 FLEX) to guide the exogenous glucose infusion rate.

*Femoral arterial blood flow* was assessed alternately in each leg, approximately 5 cm proximal to the femoral bifurcation, using an ultrasound system (Affiniti 70G; Philips). Scanning was conducted using a 12–3 MHz linear array transducer in Pulse-Wave Doppler mode. An insonation angle of <60° was maintained to ensure accurate velocity quantification. The sample volume was adjusted to encompass the entire vessel width, and the vessel diameter was manually measured as the perpendicular distance between the inner arterial walls (intima-to-intima). Blood velocity was averaged over more than five consecutive cardiac cycles, allowing the integrated software to calculate absolute blood flow. These ultrasound measurements were obtained immediately prior to each of the 13 blood sampling time points in both legs (**Figure 1A**).

*Skeletal muscle biopsies* (50–250 mg) were obtained from the vastus lateralis muscle under local anesthesia (∼3 mL of 10 mg/mL lidocaine without adrenaline [Xylocaine]; AstraZeneca) using a Bergström needle modified with manual suction. Following excision, the tissue was briefly rinsed in ice-cold saline to remove excess blood and immediately snap-frozen in liquid nitrogen. Biopsies from both the rested and exercised legs were collected at 2, 4, and 6 hours post-exercise (**Figure 2A**). For a subset of subjects (S10–S13), the 2-hour post-exercise biopsy was omitted.

*Whole-body (pulmonary) gas exchange* was determined via indirect calorimetry. Measurements were recorded for 5–10 minutes per sampling period using a canopy hood system during bed rest (Vyntus CPX Canopy; Vyaire Medical) and a face mask during the exercise bout (Vyntus CPX; Vyaire Medical). These respiratory measurements were sampled at seven distinct time points throughout the study day (**Figure 1A**).

### Biochemical analyses

#### Blood and plasma concentration

Plasma concentrations of free fatty acids (436-91995; Wako Chemicals), triacylglycerol (A11A01640; Triolab), and glycerol (GY 105; Randox Laboratories) were measured using colorimetric assays on an automated chemistry analyzer (Pentra C400; Horiba Medical). Plasma concentration of insulin (10-1113-01; Mercodia) and the catecholamines adrenaline and noradrenaline (BA E-5400; Labor Diagnostika Nord) were determined via enzyme-linked immunosorbent assay. Whole-blood concentrations of sirolimus (rapamycin) were analyzed by the Department of Clinical Biochemistry at Rigshospitalet (Copenhagen, Denmark) using liquid chromatography-tandem mass spectrometry (LC-MS/MS).

#### Tracer enrichment

Plasma enrichments of [6,6-^2^H_2_]-glucose and L-[ring-^13^C_6_]-phenylalanine were quantified using gas chromatography–mass spectrometry (GC/MSD 5977C; Agilent Technologies). Intracellular L-[ring-^13^C_6_]-phenylalanine enrichment in the skeletal muscle biopsies was measured by the Clinical Metabolomics Core Facility at Rigshospitalet, as previously described^99,100^.

#### Skeletal muscle tissue analysis

30–50 mg pieces of wet muscle tissue were freeze-dried and dissected free of visible blood, fat, and connective tissue under a stereomicroscope in a low-humidity-controlled environment. An aliquot of 6 mg (dry weight) muscle tissue was then isolated for homogenization. Ice-cold modified GSK3 buffer (pH 7.4; containing 10% [v/v] glycerol, 20 mM sodium pyrophosphate, 150 mM NaCl, 50 mM HEPES, 1% [v/v] Tergitol 15-S-40, 20 mM β-glycerophosphate, 10 mM NaF, 2 mM PMSF, 1 mM EDTA, 1 mM EGTA, 10 µg/mL aprotinin, 10 µg/mL leupeptin, 2 mM Na_3_VO_4_, and 3 mM benzamidine) was added to the freeze-dried tissue at a ratio of 100 µL/mg dry weight. Samples were homogenized at 4°C using a TissueLyser (Qiagen) with a steel bead for two 60-second cycles at 30 Hz, followed by end-over-end rotation for 60 minutes. A crude homogenate aliquot was reserved, while the remainder of the sample was centrifuged at 18,000*g* for 20 minutes at 4°C to collect the cleared lysate supernatant. Glycogen content and glycogen synthase (GS) activity were assessed directly from the crude homogenate aliquots. Glycogen levels were measured using a Fluoroskan microplate fluorometer (Thermo Fisher Scientific) following a 2-hour incubation in 2 M HCl at 95°C to hydrolyze the tissue and liberate glucosyl units. GS activity was assessed in duplicate by quantifying the generation of uridine diphosphate (UDP) in the presence of either 0.17 mM (physiological) or 8 mM (saturating) glucose-6-phosphate, as previously described^101^. Finally, the phosphorylation status of eIF4E at Ser209 was determined in the cleared lysates via standard immunoblotting procedures (antibody #9741, Cell Signaling Technology), consistent with our previously published protocol^102^.

### Calculation of physiological parameters

#### Leg glucose uptake

Calculated according to Fick’s principle by multiplying the arterial-venous blood glucose concentration difference by the femoral arterial blood flow, with the final value normalized to lean leg mass.

#### Lean leg mass

Determined by manually drawing segmented regions in the DXA analysis.

#### Whole-body glucose rate of disappearance (Rd)

Calculated using Steele’s steady-state equation^103^.

#### Mixed muscle protein synthesis

Calculated as the fractional synthesis rate (FSR) using a standard direct incorporation model^104^. Briefly, FSR was determined by calculating the change in protein-bound tracer enrichment between consecutive biopsies, relative to the average intracellular precursor enrichment over the tracer incorporation time.

#### Muscle protein breakdown

Estimated using a three-pool arteriovenous balance model across the leg utilizing mole percent excess (MPE)^104,105^. First, the inward transport of phenylalanine was determined based on venous and arterial phenylalanine concentrations, their corresponding MPE values across the intracellular, venous, and arterial pools, and femoral plasma flow. Subsequently, the rate of muscle protein breakdown (intracellular rate of appearance) was derived from the inward transport rate and the ratio of arterial to intracellular MPE, and ultimately normalized to lean leg mass.

#### Glycogen synthase activity

Expressed as fractional velocity (%FV), which was calculated by dividing the activity measured at 0.17 mM G6P by the activity measured at 8 mM G6P, as previously described^101^.

### Statistical analysis of human physiological measures

Data are presented as means ± SEM or with individual data points shown where appropriate. To evaluate the effects of the experimental interventions, two-way repeated-measures analyses of variance (2W ANOVA RMs) were performed. The specific statistical models, post hoc tests, and sample sizes for each analysis are detailed in the respective figure legends. Statistical significance was defined as P < 0.05.

### Fission yeast experiment

#### Yeast cell cultures and glucose starvation assays

Strains used in this study (Table 2). All cultures were grown at 28°C. Cells were initially inoculated in Yeast extract (YES) based media and grown overnight^106^, before being transferred into Edinburgh Minimal Medium (EMM2-N) (ForMedium, #PMD1305)^107^, supplemented with 93.5 mM NH_4_Cl (Sigma-Aldrich #09718) (EMM2). Cultures were maintained in log phase for 2 days. Before completely withdraw of glucose the cultures were exposed to low glucose. This glucose pre-conditioning step was included to extend life span of starved cells^108^. YES grown cells were transferred into EMM2 medium (ForMedium, #PMD0405) supplemented with 0.15% glucose (8.3 mM) (Sigma, #G8270) and grown for 4-5 hours before transferred into EMM2 medium (ForMedium, #PMD0405) without glucose. After 1 to 2 hours growth in EMM2 medium without glucose, approximately 300 cells were plated onto YES agar plates. Colonies formed were counted after incubation at 32 °C for a few days and defined as day 0. Samples were collected at the time points indicated in the results. Cell viability was calculated as the percentage of colony-forming units relative to day 0. Three independent experiments were performed.

**Table 1.** Subject characteristics for human *in vivo* experiment. Data presented as means ± SD. BMI, body mass index; PWL, peak work load (during one-legged kicking); VO_2_ max, maximal oxygen consumption. Mean lean leg mass is the average lean mass of the two legs.

| Subject characteristics (n=13) |  |
| --- | --- |
| Age (years) | 28 ± 4 |
| Height (m) | 1.83 ± 0.1 |
| Body mass (kg) | 74 ± 8 |
| BMI (kg/m<sup>2</sup>) | 22 ± 2 |
| Body fat (%) | 18 ± 4 |
| Lean body mass (kg) | 57 ± 7 |
| Mean lean leg mass (kg) | 10.9 ± 1.7 |
| PWL (watt) | 40 ± 9 |
| VO<sub>2</sub> max (ml/kg/min) | 50 ± 7 |

**Table 2.**
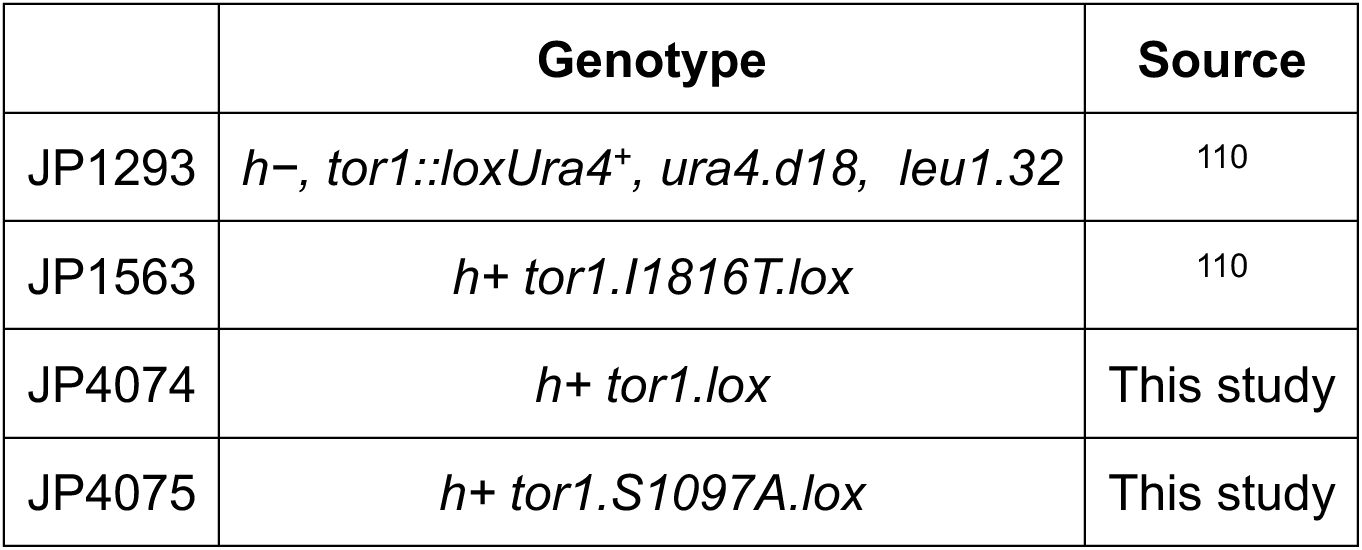
Yeast strains used in this study.

#### Generation of WT and single point mutations in tor1

WT tor1 and tor1.S1097A mutant were generated in JP1293 (*h−, tor1::loxUra4^+^, ura4.d18, leu1.32)* using cre recombinase–mediated cassette exchange (RMCE; REF^109^), as described previously^110^.

The tor1.S1097A was mutated by site-directed mutagenesis of WT tor1 in pAW8^109,110^. The recombinant plasmids carrying WT or mutant tor1 alleles were used to replace the ura4⁺ cassette in JP1293 via the Cre–lox recombination protocol^109^.

#### Statistical analysis of fission yeast experiment

Data are presented as means ± SEM. To evaluate the effects of the Tor1.S1097A mutation against WT, a 2W mixed-design ANOVA (incorporating repeated measures for the *time* factor) was used to compare the factors *mutation* vs. *time.* Sample size is n=5 with incomplete repeated measures across time. Statistical significance was defined as P < 0.05.

#### Mouse *in vivo* experimental study

All animal experiments were approved by the Danish Animal Experiments Inspectorate (license no: 2024-15-0201-01684) and complied with the European Union guidelines for the protection of vertebrate animals used for scientific purposes. Female C57BL/6J mice were acquired from Taconic Biosciences and were 12–15 weeks of age at the time of the terminal experiments. The mice were housed in a temperature-and humidity-controlled facility on a standard 12-hour light/12-hour dark cycle, with *ad libitum* access to standard rodent chow and water. To minimize circadian variation, all terminal experimental procedures were initiated at the same time of day (08:00).

#### Pilot studies

To determine the minimum effective *in vivo* dose of eFT508 required for MKNK1/2 inhibition, a total of four pilot experiments were conducted (n = 2 mice per experiment). Mice were anesthetized via an intraperitoneal injection of an FMA cocktail consisting of fentanyl (0.38 µg/g body weight; Dechra Veterinary Products), midazolam (8.0 µg/g; Accord Healthcare), and acepromazine (8.0 µg/g; Pharmaxim), administered at a volume of 10 µL/g. Body temperature was maintained at 37°C using a heating pad for the duration of the experiment. A polyethylene cannula (PE-50; Intramedic) was surgically inserted into one jugular vein, and a 5-way connector was attached to allow for the simultaneous continuous administration of anesthetics, eFT508, insulin, and glucose. Anesthesia was maintained throughout the procedure by a continuous infusion of the FMA solution (0.03 µL/g/min).

Following a 60-minute post-surgical recovery period, a 150–190 minute hyperinsulinemic-euglycemic clamp was performed, as previously described^111^. Insulin was administered as an initial bolus of 600 µU/g, followed by a constant infusion rate of 7.5 µU/g/min. Concurrently, a 20% exogenous glucose infusion was adjusted every 10 minutes based on blood glucose measurements obtained from tail vein samplings (Contour XT glucometer; Ascensia Diabetes Care) to maintain euglycemia. After 60 minutes of insulin infusion, once a steady-state glucose infusion rate (GIR) was achieved, eFT508 treatment was initiated. The eFT508 compound (0.5 µg/µL formulated in 5% dimethyl sulfoxide, 2% polysorbate 80, and 20% polyethylene glycol) was administered as an initial bolus (1, 3, or 5 µg/g), followed by a continuous infusion designed to maintain steady-state drug concentrations based on previously reported murine clearance rates (0.05 mL/min/g) and volumes of distribution (6.506 mL/g)^112^. For the specific 1 µg/g dosing cohort, an additional 10 µg/g bolus was administered after 1 hour of continuous eFT508 infusion, and the continuous infusion rate was subsequently adjusted to match the new dose.

#### Main experimental study

Based on the pilot experiments, an eFT508 dose of 3 µg/g was deemed optimal. Notably, an intraperitoneal injection of 3 µg/g was previously reported to sufficiently reduce MKNK2 signaling in the murine striatum and prefrontal cortex, whereas a 1 µg/g dose was insufficient^113^. Based on its volume of distribution, this 3 µg/g dose is estimated to yield an initial plasma concentration of ∼1.4 µM^112^.

Following catheterization surgery, a 3 µg/g bolus of eFT508 or an equivalent volume of vehicle was administered over a 50-minute period to minimize the acute injection rate. This initial bolus was followed by a continuous maintenance infusion (0.023 µg/g/min) for the remainder of the experiment. Anesthesia, insulin and glucose infusions, and blood glucose monitoring were performed as described for the pilot experiments. Eighty minutes into the hyperinsulinemic-euglycemic clamp, a 25 µCi bolus of [^3^H]-2-deoxyglucose (2-DG) in 65 µL of isotonic saline was administered intravenously. A terminal blood sample was collected at 90 minutes to determine plasma specific radioactivity and confirm similar insulin exposure (**Figure S3C**). Immediately thereafter, mice were euthanized via cervical dislocation. Skeletal muscle (tibialis anterior, quadriceps, and gastrocnemius), perigonadal white adipose tissue, interscapular brown adipose tissue, heart, and brain tissues were rapidly excised and snap-frozen in liquid nitrogen.

The study initially included *n* = 6 mice per treatment group. However, one mouse in the vehicle group died during the insulin clamp procedure and was consequently excluded from the dataset, resulting in a final sample size of *n* = 6 for the eFT508 group and *n* = 5 for the vehicle group.

### Biochemical analysis

#### Tissue homogenization

Tissue was ground into powder under liquid nitrogen using a mortar and pestle, from which 20–40 mg was aliquoted for subsequent homogenization. Ice-cold modified GSK3 buffer was added to the frozen tissue at a ratio of 10–20 µL/mg. Homogenization was performed at 4°C through a multi-step procedure: two 60-second cycles of ultrafast shaking at 30 Hz in a TissueLyser (Qiagen) with a steel bead, followed by probe sonication at 20% output for 20 seconds, end-over-end rotation for 60 minutes, and a final centrifugation at 5,000*g* for 5 minutes. The resulting cleared supernatant was collected as the sample lysate.

#### Immunoblotting

Performed on the sample lysates using the procedures described above. The primary phospho-specific antibodies (all obtained from Cell Signaling Technology) targeted the following proteins and residues: eIF4E at Ser209 (#9741), Akt at Ser473 (#9271), PRAS40 at Thr246 (#2997), p70 S6K at Thr389 (#9205), TBC1D4 at Thr649 (#8881), AMPKα2 at Thr172 (#2531), p38 MAPK at Thr180/Tyr182 (#9211), and 4E-BP1 at Thr37/46 (#9459).

#### Tissue [³H]-2-DG uptake

Determined by quantifying the intracellular accumulation of phosphorylated [^3^H]-2-deoxyglucose ([^3^H]-2-DG-6-P). Briefly, two 100 µL aliquots of the tissue lysate were processed as previously described^114^. Circulating [^3^H]-2-DG specific activity was determined from 5 µL of plasma. All processed tissue and plasma samples were analyzed using a liquid scintillation counter (PerkinElmer) with 3 mL of Ultima Gold scintillation fluid.

#### Plasma insulin concentration

Determined via a human insulin specific enzyme-linked immunosorbent assay (10-1113-01; Mercodia).

### Calculation of 2-deoxyglucose uptake

Because the full plasma decay curve could not be sampled without disturbing the steady state of the clamp, the AUC for ^3^H-2-DG was modelled using a mono-exponential decay function validated by reference sampling. The theoretical initial tracer concentration (C_0_) at t = 0 min was calculated for each animal based on the injected dose normalized to an assumed glucose distribution volume of 200 mL/kg^115,116^. The individual elimination rate constant (k) was then derived for each mouse by calculating the slope required to connect the theoretical C_0_ to the measured plasma radioactivity at t=10 min, using the formula:

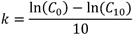

Total plasma tracer exposure (AUC) was obtained by integrating the decay curve from tracer administration (t = 0 min) to the time of tissue collection (t = 11 min):

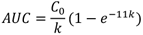

Tissue ^3^H-2-DG uptake (*R_g_*) was subsequently determined by dividing the accumulated tissue radioactivity by the integrated plasma tracer exposure:

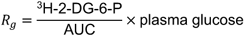

### Statistical analysis of mouse physiological measures

Data are presented as means ± SEM, with individual data points shown where appropriate. To evaluate the physiological effects of eFT508 compared to vehicle, unpaired Student’s *t*-tests were utilized for comparisons between two independent groups. To evaluate the effect of eFT508 on blood glucose following drug administration, a 2W mixed-design ANOVA (incorporating repeated measures for the *time* factor) was used to compare the factors *eFT508* vs. *time.* The specific statistical settings, post hoc tests, and sample sizes for each analysis are detailed in the respective figure legends. Statistical significance was defined as P < 0.05.

### Human *ex vivo* experimental study

This *ex vivo* study was approved by the Committee on Health Research Ethics of the Capital Region of Denmark (Reference number: H-25032376) and performed in accordance with the ethical standards laid down in the Declaration of Helsinki. All subjects provided written informed consent after receiving comprehensive oral and written information regarding the study procedures.

### Screening

Six healthy male subjects (aged 22–35 years with no known underlying diseases) were initially screened via telephone interviews. All six individuals met the inclusion criteria, participated in the clinical screening, and subsequently completed the entire experimental protocol.

On the clinical screening day, subjects were instructed to consume a substantial breakfast, while strictly abstaining from coffee, two hours prior to arriving at the laboratory. Following 30 minutes of supine rest, an antecubital venous catheter was inserted. A preliminary blood sample was drawn and immediately analyzed to confirm normal baseline levels of hemoglobin and blood glucose. Subsequently, 400 mL of whole blood was collected into multiple evacuated serum separator tubes (SST) containing a clot activator. The blood was allowed to clot at room temperature for 30 minutes. The tubes were then centrifuged at 1,300*g* for 10 minutes, and the resulting serum was collected, aliquoted, and stored at -30°C for use in the subsequent *ex vivo* tissue incubations.

### Experimental design

Subjects were instructed to refrain from physical activity and alcohol consumption for three days prior to the experimental day. Upon arrival at the laboratory, subjects initially rested in a hospital bed for one hour. Subsequently, three successive muscle biopsies were obtained at 30-minute intervals using the procedures described above. Immediately following excision, the biopsies were briefly rinsed in room-temperature (RT) saline, submerged in pre-oxygenated RT Krebs-Ringer buffer (pH 7.3; containing 117 mM NaCl, 4.7 mM KCl, 2.5 mM CaCl_2_, 1.2 mM KH_2_PO_4_, 1.2 mM MgSO_4_, 24.6 mM NaHCO_3_, 5 mM glucose, and 5 mM mannitol), and rapidly transported to the *ex vivo* laboratory (∼1 min).

Upon arrival at the laboratory, each biopsy was meticulously dissected into four longitudinal muscle fiber bundles (approximately 5–10 mm in length and 5–20 mg wet weight). To preserve biomechanical tension, the bundles were mounted with fine needles on a customized silicone block. The mounted tissue was incubated in 100% human serum at 30°C and continuously gassed with a 95% O_2_ / 5% CO_2_ mixture to maintain a strict physiological pH of 7.4. To eliminate inter-day and inter-subject media variance, the incubation serum consisted of evenly mixed, pooled serum aliquots obtained from all study participants during the screening visit.

The fiber bundles were incubated for a total duration of 60 minutes in the presence of either 1 µM eFT508 or a vehicle control (DMSO; 0.1% final volume). For the final 30 minutes of the incubation period, insulin (10,000 µU/mL) was added to half of the fiber bundles. Following the 60-minute incubation, the fiber bundles were rapidly dismounted, rinsed in ice-cold saline, and snap-frozen in liquid nitrogen. The optimal media concentrations of eFT508 and insulin utilized in this protocol were determined during preliminary dose-response experiments conducted for this study (**Figures S4A** and **S4B**).

### Biochemical analysis

*Tissue lysis and immunoblotting:* On the experimental study day, three distinct muscle fiber bundles were incubated under each specific treatment condition. Following incubation, these bundles were individually lysed in 100 µL of SDC lysis buffer (detailed below) and subsequently pooled to minimize intra-condition variance. A small aliquot of this pooled lysate was mixed with Laemmli buffer (final concentrations: 53 mM Tris-HCl [pH 6.8], 91 mM DTT, 52.5 mM SDS, 4.5% glycerol, and 27 µM bromophenol blue) and subjected to immunoblotting as described above. Specifically, the samples were probed for the phosphorylation of eIF4E at Ser209, Akt at Ser473, and PRAS40 at Thr246. This initial immunoblotting step successfully verified the targeted pharmacological inhibition of MKNK2 signaling (**Figure S4C**) and the robust induction of the canonical insulin signaling cascade (**Figures S4D** and **S4E**). The remainder of the pooled lysates was reserved and further processed for phosphoproteomic analysis (detailed below).

### Phosphoproteomics

#### Preparation of samples for phosphoproteomics

Frozen muscle biopsies (10–20 mg wet weight) were ground into a fine powder under liquid nitrogen using a mortar and pestle and immediately lysed in 200 µL of SDC lysis buffer (4% sodium deoxycholate, 100 mM Tris, pH 8.5) at 95°C for 5 minutes with agitation at 2,000 rpm. After cooling to RT, the samples were sonicated using a tip probe sonicator (SFX150; Branson) for a total of 30 seconds (3-second on/3-second off pulses at 30% amplitude). Lysates were subsequently centrifuged at 14,000 × *g* for 15 minutes at RT. The cleared supernatants were collected, and protein concentrations were determined using a bicinchoninic acid (BCA) protein assay (Pierce; Thermo Fisher Scientific). A total of 700 µg of protein per sample was simultaneously reduced and alkylated with 1 mM tris(hydroxypropyl)phosphine (THPP) and 40 mM chloroacetamide (CAA) by incubating at 45°C for 5 minutes, followed by 40 minutes at RT. Proteins were then sequentially precipitated using a chloroform/methanol extraction followed by an acetone precipitation. The resulting purified protein pellets were resuspended in SDC lysis buffer, and a final 200 µg aliquot of protein from each sample was processed for phosphopeptide enrichment utilizing the EasyPhos workflow^117^.

#### Liquid-chromatography-tandem mass spectrometry

Enriched phosphopeptides in MS loading buffer (2% ACN, 0.3% TFA) were loaded onto in-house fabricated 55-cm columns with a 75-µm I.D. and packed with 1.9 µm C18 ReproSil Pur AQ particles using a Vanquish Neo coupled to an Orbitrap Astral mass spectrometer (Thermo Fisher Scientific). Column temperature was maintained at 60 °C using a Sonation column oven, and peptides separated using a binary buffer system comprising 0.1% formic acid (buffer A) and 80% ACN plus 0.1% formic (buffer B), at a flow rate of 400 nL/min with a gradient of 3-19% buffer B over 40 min followed by 19-41% buffer B over 20 min, resulting in ∼60 min gradients. Peptides were analyzed with one full scan (380-980 m/z, R = 240,000) at a target of 5,000,000 ions, followed by 300 data-independent acquisition (DIA) MS/MS scans (380-980 m/z) with HCD (target 50,000 ions, maximum injection time 3.5 ms, isolation window 2 m/z, NCE 24%), with fragments detected in the Astral mass analyzer.

#### Raw MS data processing

##### In vivo

Raw data were analysed using direct DIA analysis in the Spectronaut software (version 18.6.231227.55695). A spectral library was generated against the Human UniProt FASTA database (accessed in January 2024). Default settings were used for search parameters. The following preprocessing and analyses were performed in the R software environment version 4.5.2. To remove unreliable phosphosites, we first removed all phosphosites with a maximum localisation probability < 0.75 (retaining only class I phosphosites). Raw intensities were log2 transformed and phosphosites with log2 values < 1 were converted to missing values as they were considered unreliable measurements indistinguishable from background noise. We then performed centre-median normalisation followed by the addition of a constant offset of 10.0 to restore positive log_2_ intensity scaling.

##### Ex vivo

Raw data were analysed using direct DIA analysis in the Spectronaut software (version 19.3.241023.62635). A spectral library was generated against the Human UniProt FASTA database (accessed in March 2025). Default settings were used for search parameters. The following preprocessing and analyses were performed in the R software environment version 4.5.2. To remove unreliable phosphosites, we first removed all phosphosites that had a localisation probability of < 0.5, then any remaining phosphosites with a maximum localisation probability < 0.75 were further removed (retaining only class I phosphosites). Raw intensities were log2 transformed and phosphosites with log2 values < 3 were converted to missing values as they were considered unreliable measurements indistinguishable from background noise. We then performed centre-median normalisation followed by the addition of a constant offset of 10.0 to restore positive log_2_ intensity scaling.

#### Quality control and data filtering

##### Human in vivo dataset

For quality control of the *in vivo* human study, phosphosites were initially filtered to retain those present in >70% of all samples. Duplicate phosphopeptides sharing identical quantification due to multiple phosphorylation were collapsed into a single representative value. Next, a correlation matrix based on Pearson’s correlation coefficients for all pairwise observations was generated. This matrix identified 7 of the 104 samples as potential outliers (**Figure S2A**). However, a *t*-distributed stochastic neighbor embedding (*t*-SNE) plot demonstrated that the samples clustered primarily by subject ID (**Figure S2B**), consistent with previous observations^23^, with no clear systematic pattern associated with the seven outlier samples identified in the correlation matrix. Similar observations were obtained with principal component analysis (PCA). Therefore, no samples were excluded from the final *in vivo* dataset.

##### Human ex vivo dataset

For quality control of the *ex vivo* human study, phosphosites were filtered to retain those present in >60% of all samples. Duplicate phosphopeptides sharing identical quantification due to multiple phosphorylation were collapsed into a single representative value. A Pearson correlation matrix revealed that 3 of the 24 samples clustered as slight potential outliers (S3 Insulin DMSO, S4 Basal DMSO, and S4 Basal eFT508; **Figure S4F**). While a *t*-SNE plot showed sample clustering by subject ID without apparent outliers (**Figure S4G**), PCA indicated that these same three samples separated distinctly along principal components 2 and 3 (PC2 and PC3; **Figures S4H+I**). Hierarchical clustering of *z*-scored samples demonstrated that the two S4 Basal samples were the most distinctive from the rest of the dataset (**Figures S4J+K**). Consequently, the two S4 Basal samples were excluded from the final *ex vivo* dataset.

#### Bioinformatics analysis of phosphoproteomics data

##### Human in vivo study

A rigorous three-step statistical analysis was performed to identify phosphorylation sites associated with rapamycin-induced changes in insulin sensitivity (**Figure 3A**). First, four biopsy conditions of interest were defined, as detailed in the *Results* section. For each condition, a paired linear model was performed to test the effect of rapamycin versus placebo. Sites exhibiting a P < 0.05 and an absolute fold change > 1.5 were considered significant. Next, these significant sites were evaluated for the main effects of insulin, exercise, or their interaction using a paired multilevel linear model. In this secondary analysis, the treatment condition (rapamycin versus placebo) was collapsed, meaning subject data from the rapamycin and placebo trial days were treated as independent biological replicates. This approach increases statistical power for detecting exercise and insulin effects by doubling the number of observations, but assumes that the exercise and insulin responses are qualitatively similar across drug conditions. We consider this assumption reasonable given that the sites were pre-filtered for a rapamycin effect in step 1, and step 2 tests an orthogonal biological question (physiological regulation irrespective of drug treatment). Because these sites were already pre-filtered for the rapamycin effect, an unadjusted P < 0.05 was considered significant for this step. Finally, sites passing the first two analytical thresholds were tested for their association with the rapamycin-induced change in insulin sensitivity. Specifically, subject-paired phosphorylation levels between the rapamycin and placebo days were correlated with the corresponding exercise-induced change in the area under the curve (AUC) for insulin-stimulated glucose uptake (**Figure 1C**) using repeated-measures correlation analysis^118^. Notably, sites were only correlated within the specific biopsy condition in which they were initially found to be significantly regulated by rapamycin. Statistical significance for this final step was defined as P < 0.05 combined with a bootstrap-determined 95% confidence interval (CI) excluding zero.

Unadjusted p-value thresholds were applied at each step of the pipeline. This analytical approach reflects the relatively modest perturbation of the phosphoproteome achievable at the maximum safe oral dose of rapamycin in humans (0.21 mg/kg; affecting 2-4% of quantified sites), where stringent correction at any single step risks eliminating biologically meaningful signals. The sequential application of three filtering criteria, each addressing a distinct biological question (drug response, physiological regulation, and phenotypic association), provides cumulative selectivity while preserving sensitivity to novel biology. We note that filters 1 and 3 are not statistically independent, as both reflect the rapamycin perturbation within the same individuals; rather, filter 3 tests whether the inter-individual variance in the drug effect on phosphorylation tracks with the variance in the physiological outcome. Filter 2 (exercise and insulin regulation) provides the independent biological constraint. The 36 sites identified at the terminal step should nonetheless be considered hypothesis-generating candidates, and the key sites were subsequently validated through pharmacological intervention in independent model systems. For all sites significantly regulated by rapamycin versus placebo, the delta (Δ) phosphorylation level was calculated within the specific biopsy condition. To standardize the directionality of the biological response, sites with a median Δ value < 0 were mathematically inverted (multiplied by -1) prior to z-score normalization across the cohort. This transformation ensured that the strongest responder to rapamycin for any given site consistently possessed the highest z-score, regardless of whether the original site was upregulated or downregulated by the drug. These z-scored, rapamycin-regulated sites were subsequently hierarchically clustered to visualize the inter-individual diversity among top responders, and the subject-specific distributions of z-scores were quantified for visualization. Furthermore, to identify clusters of co-regulated sites, a correlation matrix based on repeated-measures correlation was generated for all sites that passed the filtering criteria for correlation with rapamycin-induced changes in insulin sensitivity.

Lastly, a select subset of functionally relevant sites (MKNK2 S74, EIF4G1 S1209, and EIF4G1 S1124) was further assessed for their association with the exercise-induced change in insulin sensitivity on the placebo day. Specifically, the specific phosphorylation levels in the rested (REST) and previously exercised (PEX) legs during insulin stimulation on the placebo day were correlated with the corresponding steady-state (mean 80–120 minutes) insulin-stimulated glucose uptake using repeated-measures correlation, consistent with our previous methodology^23^.

##### Human ex vivo study

The pharmacological effect of eFT508 was evaluated across three specific conditions of interest: the basal state, the insulin-stimulated state, and a combined state (to maximize statistical power). For each condition, a paired linear model was utilized to determine the effect of eFT508 versus the DMSO control. Sites exhibiting a P < 0.05 and an absolute fold change > 1.5 were considered statistically significant.

### R packages and analytical settings

All linear models were executed using the *limma* package^119^ applying empirical Bayes moderation. Repeated-measures correlations were performed using the *rmcorr* package^118^, with bootstrapping utilized to calculate confidence intervals. All heatmap visualizations were generated via the *ComplexHeatmap* package^120^. For the clustering of z-scored rapamycin-regulated sites, hierarchical clustering was applied using Pearson correlation as the distance metric alongside Ward’s minimum variance method (ward.D2). For the repeated-measures correlation matrix, hierarchical clustering was performed using Euclidean distance and complete linkage.

## Data visualization

Statistical graphs and data plots were generated using GraphPad Prism (version 11.0.0) or RStudio (utilizing the *ComplexHeatmap* package^120^). Final figure assemblies and aesthetic refinements were subsequently performed in Adobe Illustrator (version 29.1). Study design schematics and intracellular signaling pathway diagrams were initially created using BioRender (BioRender.com) and finalized in Microsoft PowerPoint (version 16.106).

## Acknowledgement

We would like to thank the following colleagues for their contribution to the study: Elise J. Needham (University of Cambridge), Janne R. Hingst (University of Copenhagen (UCPH), currently Novo Nordisk inc.), Jørgen Jensen (Norwegian School of Sport Sciences), and Lars Holm (UCPH) for scientific discussions; Le Lene S. Stevner (UCPH) for support on data regulation and protection; Lina H. H. Le (Murdoch Children’s Research Institute (MCRI)), James Burgess (MCRI), Irene B. Nielsen (UCPH) and Betina Bolmgren (UCPH) for their skilled technical help in the laboratory; Erik A. Richter (UCPH) and Erik K. A. Niklasson (UCPH) for clinical support during human experimentation; Gerrit v. Hall at the Clinical Integrative Fluxomics core (UCPH) for sample analysis; Jonas R. Larsen at the Mechanical Workshop (UCPH) for manufacturing of experimental equipment.

M.R.L. and J.D.O. were supported by a research grant from the Danish Diabetes and Endocrinology Academy, which is funded by the Novo Nordisk Foundation, grant no. NNF17SA0031406. J.F.P.W. was supported by The Novo Nordisk Foundation (NNF 1235861001 and NNF1204611001). D.E.J is an Australian Research Council Laureate Fellow (FL200100096). S.J.H. is supported by a National Health and Medical Research Council (NHMRC) Investigator Grant (2026905), Synergy Grant (2035975), and is a member of The Novo Nordisk Foundation Center for Stem Cell Medicine, reNEW, which is supported by Novo Nordisk Foundation grant no. NNF21CC0073729. J.P. and S.J.H. were supported by Australian Research Council [DP220103531]. R.K. was supported by an EFSD and Novo Nordisk Foundation Future Leaders Award (NNFSA250106138).

## Author contribution

Conceptualization, M.R.L., D.E.J., R.K., S.J.H., J.F.P.W.; Human *in vivo* experiment, M.R.L., J.D.O., N.S.H., J.M.K., M.N.Z., and J.F.P.W.; Yeast experiment, T.W. and J.P.; Mouse experiment, M.R.L. and K.A.S.; Human *ex vivo* experiment, M.R.L., J.D.O., M.N.Z., R.E., J.T.E.M., J.S.J., J.F.P.W.; Biochemical analyses, M.R.L., J.B.B., E.S.S.; Bioinformatic analyses, M.R.L. and H.H.; Supervision, R.K., S.J.H., and J.F.P.W.; Funding acquisition, M.R.L., J.D.O., J.R.G., D.E.J., S.J.H., and J.F.P.W.; Visualization, M.R.L.; Writing original draft, M.R.L., H.H., S.J.H., and J.F.P.W.; Review and editing, all authors.

---

**Figure S1.**
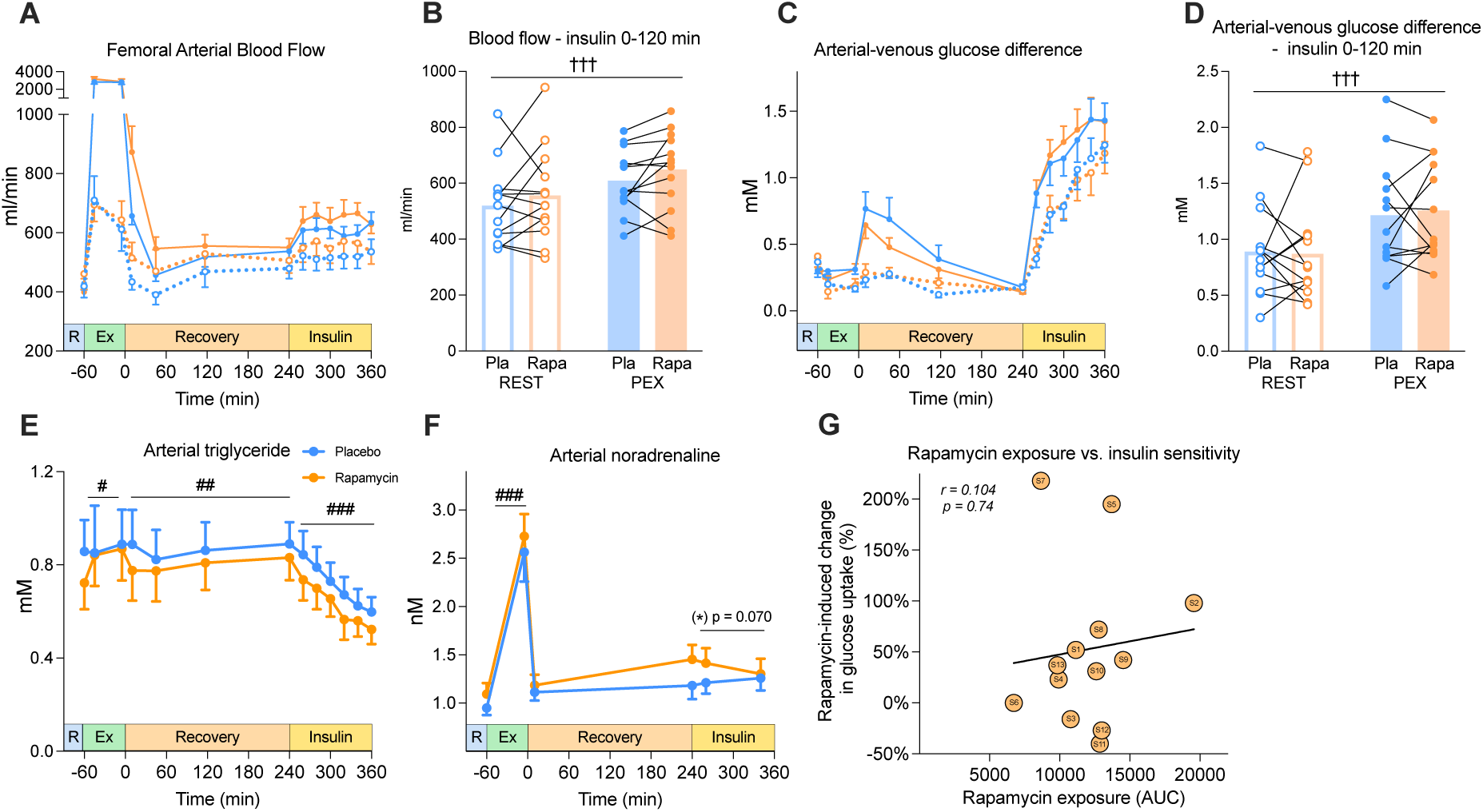
Rapamycin modulates whole-body metabolism and skeletal muscle insulin sensitivity. **A**) Femoral arterial blood flow throughout the study protocol (n=9 for the first 6 time points). **B**) Mean femoral arterial blood flow during insulin clamp (0-120 min). **C**) Arterial-venous (AV) glucose difference throughout the study protocol (n=9 for the first 6 time points). **D**) Mean AV glucose difference during insulin clamp (0-120 min). **E**) Arterial plasma triglyceride and **F**) noradrenaline concentrations throughout the study protocol (n=9 for the first 6 time points). **G**) Correlation between the effect of rapamycin on exercise-induced change in insulin sensitivity (from Figure 1E) and area under the curve (AUC) for whole-blood rapamycin concentration across the study day (from Figure 1F). Data in **[A, C, E–F]** are means ± SEM; data in **[B, D]** are mean with individual values paired (*placebo* vs. *rapamycin*). Two-way repeated measures ANOVA was used to test the following factors: *rapamycin* vs. *exercise* **[B, D]**; *time* vs. *rapamycin* within each condition (exercise, recovery, or insulin), using the final time point of the preceding condition as baseline **[E–F]**. Pearson correlation was used to evaluate association between change in insulin sensitivity and rapamycin exposure [**G**]. Straight lines indicate a main effect. ††† p<0.001 effect of *exercise*; (*) p = 0.07 (borderline); effect of *rapamycin*; ### p<0.001, ## p<0.01, # p<0.05 effect of *time*.

**Figure S2.**
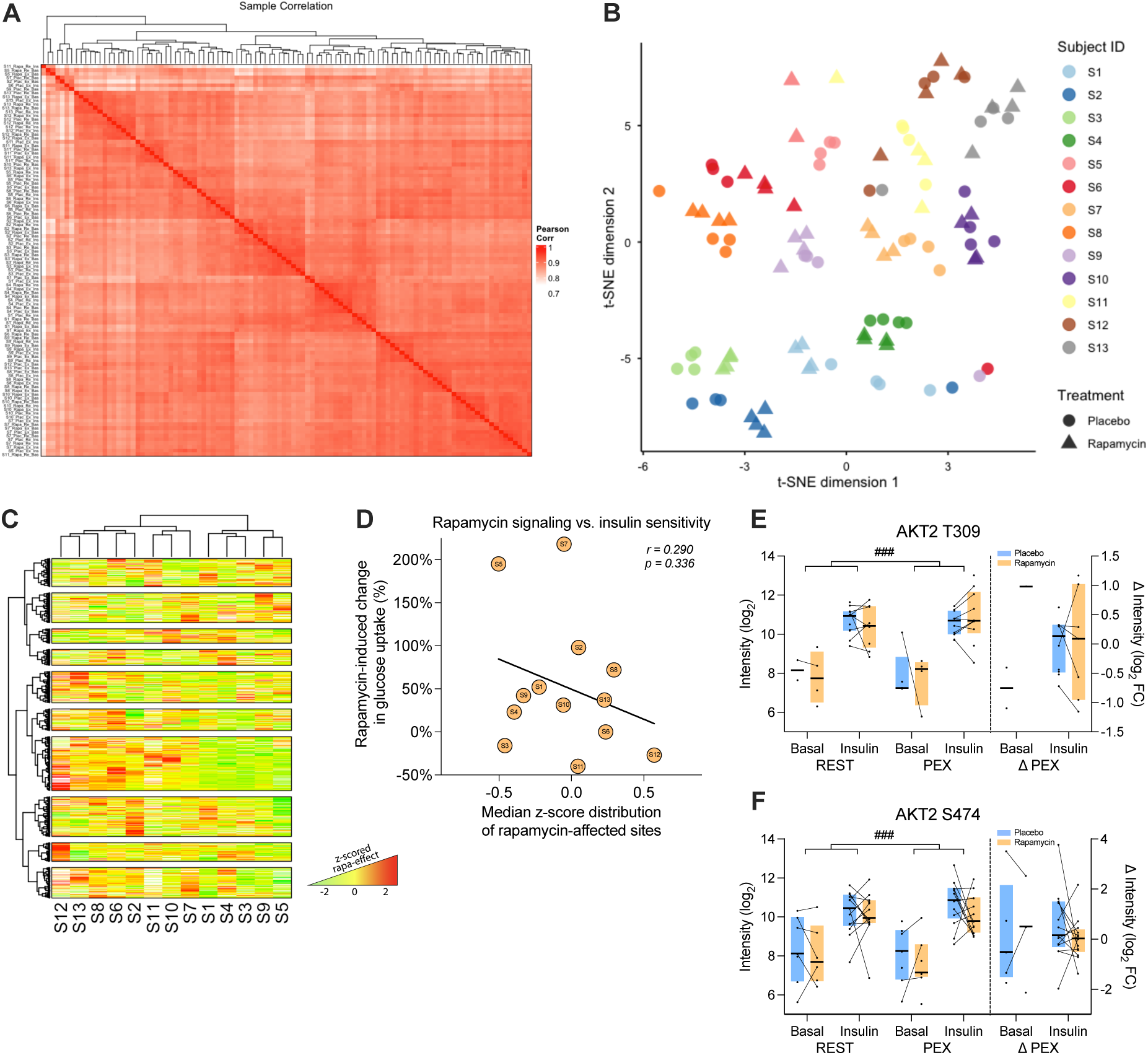
Personalized phosphoproteomics identify a subset of sites linked to rapamycin-induced improvements in muscle insulin sensitivity following exercise. **A**) Pearson correlation matrix of the phosphoproteomes. **B**) Sample phosphoproteome clustering by t-SNE. **C**) Subject-specific effects of rapamycin relative to the group mean for all rapamycin-responsive sites (from 3B). For visual consistency, sites with a median < 0 were inverted (multiplied by -1), and all values were z-score normalized across the cohort (i.e., the greatest rapamycin response corresponds to the highest z-score). **D**) Correlation between the effect of rapamycin on exercise-induced change in insulin sensitivity (from Figure 1E) and the medians of z-scored rapamycin-affected phosphosites (from Figure 3C). **E**) Phosphorylation levels of AKT2 T309, and **F**) AKT2 S474. Data in [**E+F**] are presented as medians ± interquartile range. Statistical analyses included: Pearson correlation [**D**], linear models [**E+F**] to test the effect of drug treatment (*rapamycin* vs *placebo*), and two-way repeated measures ANOVA [**E+F**] to test the factors *exercise* vs. *insulin*, independent of rapamycin treatment. Straight lines indicate a main effect. ### p<0.001 effect of *insulin*.

**Figure S3.**
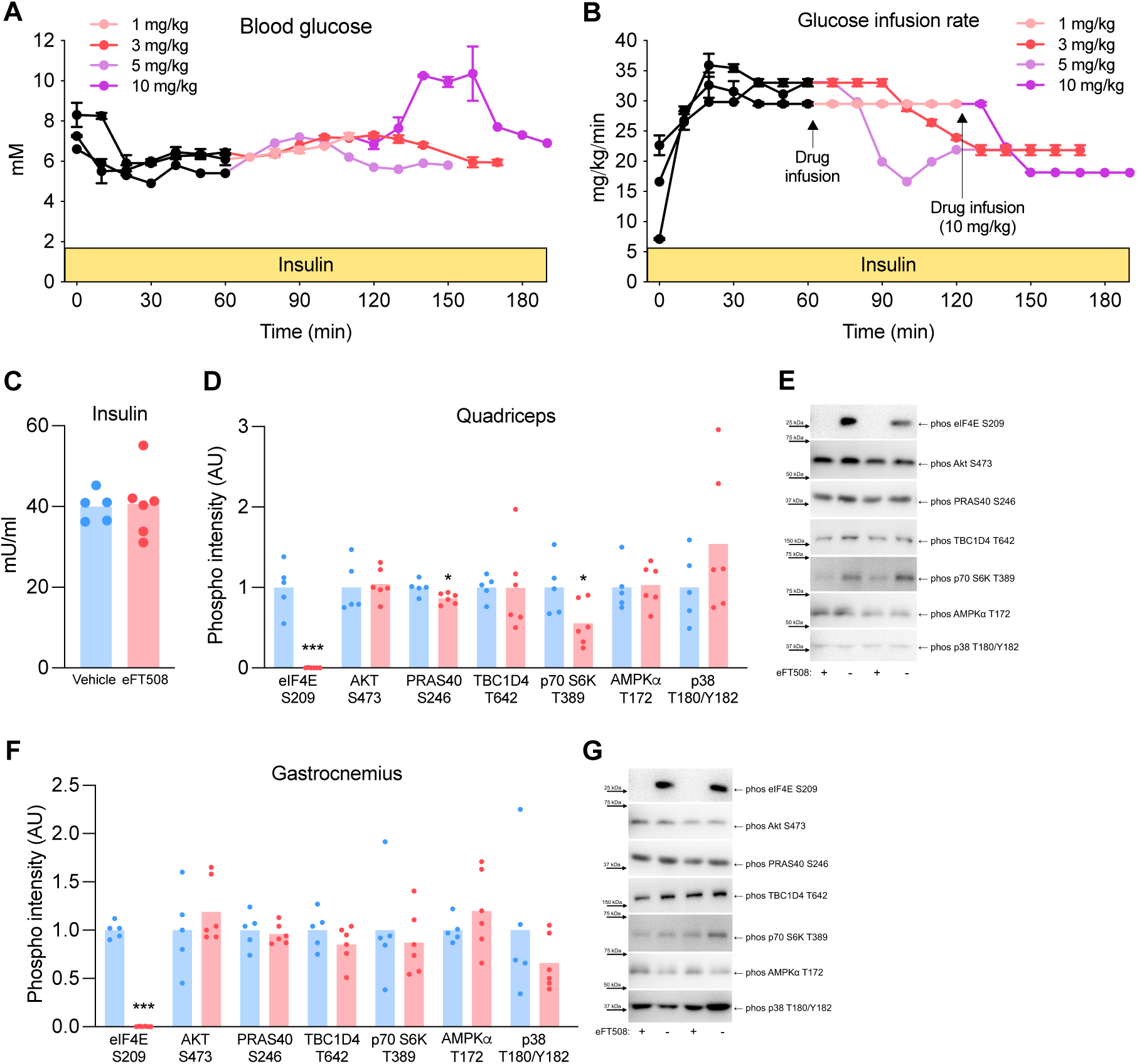
Pharmacological inhibition of MKNK2 reduces muscle insulin sensitivity. **A)** Tail-vein blood glucose concentrations during the pilot study protocol (n=1-2). Transition from black to colored lines indicates the initiation of intravenous drug infusion at the specified concentrations. **B)** Glucose infusion rate (GIR) during the hyperinsulinemic-euglycemic clamp. **C**) End-stage plasma insulin concentration. Immunoblotting of selective phosphorylation sites in **D**) *quadriceps* muscle, with **E**) corresponding representative immunoblots, and **F**) *gastrocnemius* muscle, with **G**) corresponding representative immunoblots. Data in **[A-B]** are means ± SEM; data in **[C-D+F]** are mean with individual values. Unpaired t-tests were used to compare *eFT508* vs. *vehicle* **[C-D+F]**. *** p<0.001, ** p<0.01, * p<0.05 effect of *eFT508*.

**Figure S4.**
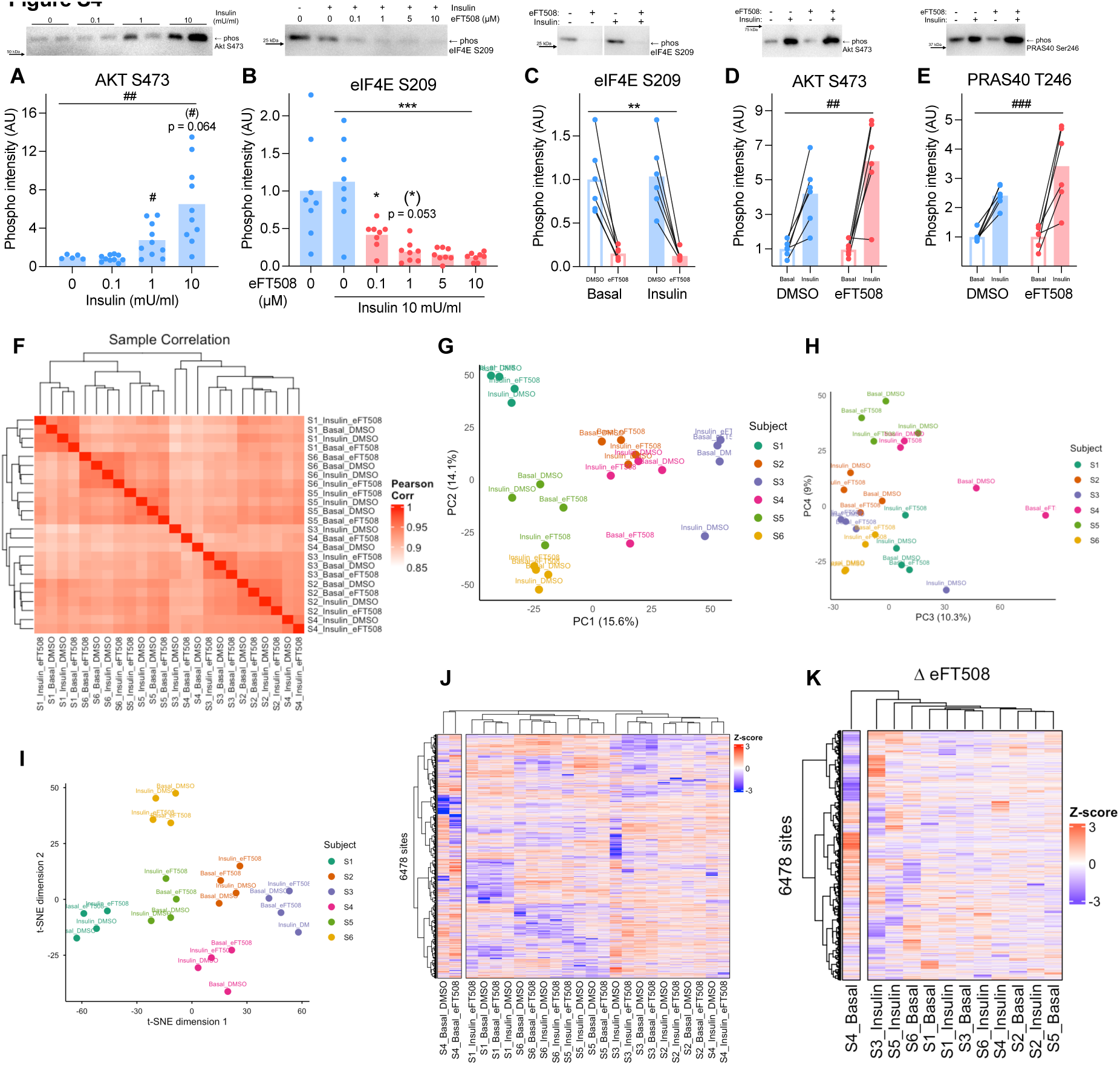
Delineating the MKNK2 signaling axis in human skeletal muscle. **A**) Immunoblotting of AKT S473 in response to increasing insulin concentrations during *ex vivo* incubation. **B**) Immunoblotting of eIF4E S209 with increasing concentration of eFT508 during ex vivo incubation of human skeletal muscle. **C**) Immunoblotting of eIF4E S209, **D**) AKT S473 and **E**) AKT1S1 S246 in *ex vivo* incubated human skeletal muscle samples (from Figure 6B). **F**) Spearman correlation matrix of the global phosphoproteomes (samples from Figure 6B). **G**) Principal component analysis (PCA) of phosphoproteome clustering by PC1 and PC2, and **H)** PC3 and PC4. **I)** t-Distributed Stochastic Neighbor Embedding (t-SNE) visualization of phosphoproteomic profiles. **J)** Hierarchical heatmap clustering of raw phosphoproteomic data, and **K)** treatment-induced changes expressed as eFT508 (eFT508 – DMSO). Data in **[A-E]** are presented as means with individual values. Lines connect samples from the same subject across drug condition (*DMSO* vs. *eFT508*) or insulin stimulation (*Basal* vs. *Insulin*). To test the effect of increasing dosis of insulin or eFT508 one-way Welch’s ANOVAs were used, which accounted for variance heterogeneity across doses **[A-B]**; Two-way repeated measures ANOVA was used to test the factors *basal/insulin* vs. *DMSO/eFT508* **[C-E]**. Main effects are indicated by straight lines. Significant interactions (p<0.05) were followed by Dunnett’s T3 multiple comparison tests, by selectively testing if a specific dose was different from the prior. # p<0.05, ## p<0.01, ### p<0.001 effect of *insulin*; *** p<0.001, ** p<0.01, * p<0.05 effect of *eFT508*.

**Figure S5.**
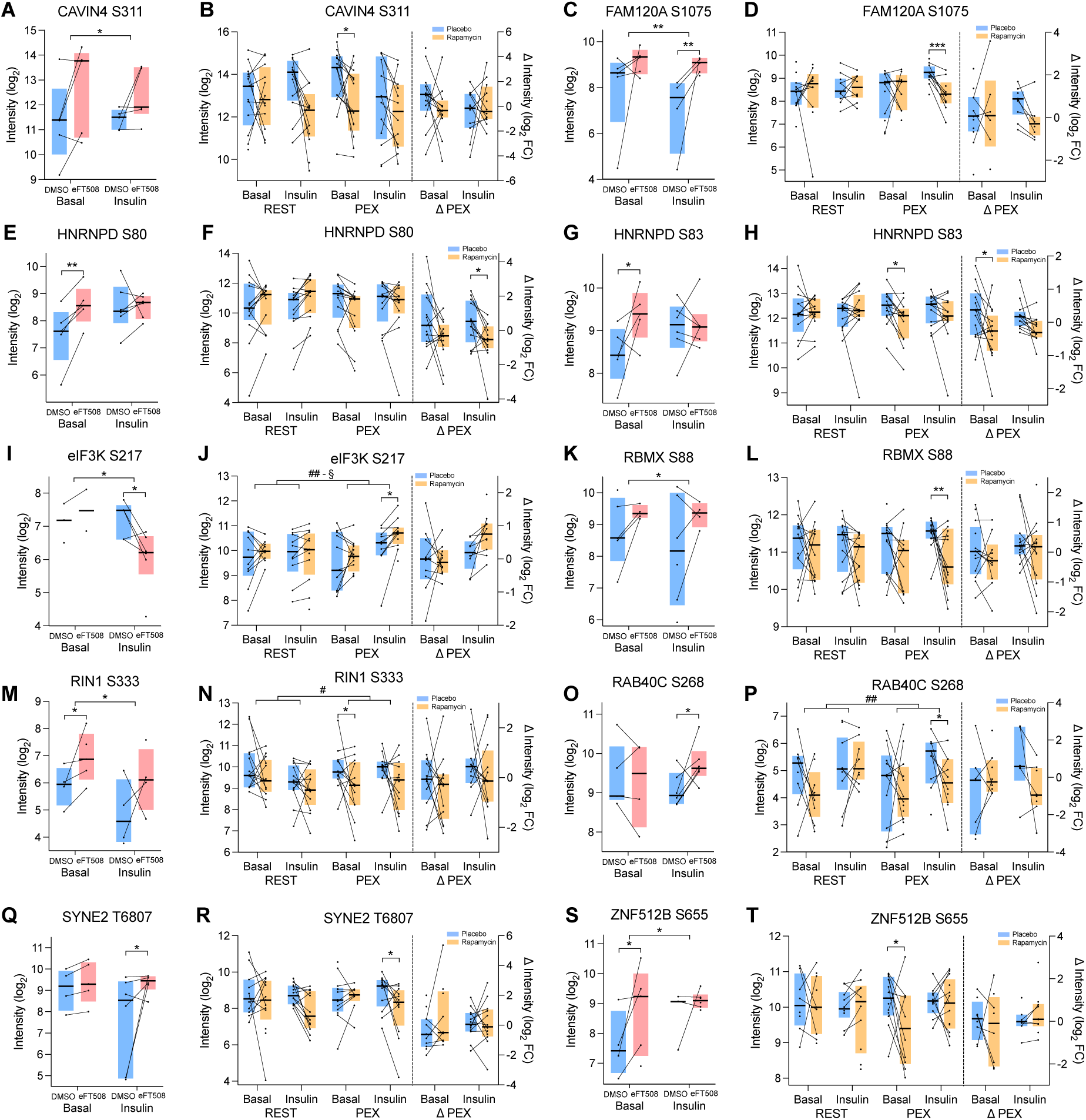
**Delineating the MKNK2 signaling axis in human skeletal muscle. A-T**) Phosphorylation levels of selective sites displaying divergent regulation by eFT508 in *ex vivo* incubated human muscle and in muscle samples from subjects treated with rapamycin *in vivo*. Data in **[A-T]** are presented as median ± interquartile range. Lines connect samples from the same subject across drug condition (*placebo/DMSO* vs. *rapamycin/eFT508*) [**A-T**]. Statistical analyses included: paired linear models [**A-T**] to test the effect of drug treatment (*rapamycin/eFT508* vs *placebo/DMSO*); two-way repeated measures ANOVA to test the factors *exercise* vs. *insulin*, independent of rapamycin treatment **[B, D, F, H, J, L, N, P, R, T]**. Main effects are indicated by straight lines. Significant interactions (p<0.05) were followed by Tukey’s post hoc tests. § p<0.05 *interaction* of factors; *** p<0.001, ** p<0.01, * p<0.05 effect of *rapamycin* or *eFT508*; # p<0.05 effect of *insulin*.

